# Cell-envelope conduits enable transfer of megabase-sized double-stranded DNA between cells of the nosocomial pathogen *Acinetobacter baumannii*

**DOI:** 10.64898/2026.09.02.748809

**Authors:** Sina Manger, Clara Börnsen, Sara Garcia Torres, Pauline Roth, Julian Sommer, Ashwin Balakrishnan, Felix Langschied, Celina Thiel, Inga Hänelt, Mike Heilemann, Stephan Göttig, Ingo Ebersberger, Klaas M. Pos, Achilleas S. Frangakis

## Abstract

Horizontal gene transfer in *Acinetobacter baumannii* has been attributed to conjugative pili, transduction, and natural transformation. Here, we describe a previously unrecognized type of chromosomal DNA exchange mediated by cell-envelope conduits that establish direct cytoplasmic continuity between neighboring *A. baumannii* cells. Cryo-electron tomography reveals that these conduits, which were found in multiple clinical isolates, comprise an outer membrane, a peptidoglycan layer, and an inner membrane. The ∼65 nm-thick conduits contain two ∼2 nm-thick filaments, which super-resolution microscopy identifies as DNA. Interstrain horizontal gene transfer assays coupled with whole-genome sequencing demonstrate the transfer and homologous recombination of chromosomal segments up to 1.1 Mbp, corresponding to as much as 27% of the *A. baumannii* genome. Together, these findings provide direct structural and functional evidence for a conduit-mediated large-scale chromosomal DNA exchange, thus expanding the known repertoire of horizontal gene transfer strategies that may contribute to the remarkable genomic plasticity of *A. baumannii*.

## Introduction

*Acinetobacter baumannii* is a highly successful opportunistic pathogen that causes severe infections, including pneumonia and wound and bloodstream infections, and is associated with high mortality rates ^1,2^. Due to its remarkable adaptability, internationally disseminated clonal lineages (IC) persist across a wide range of environments, including healthcare settings ^2–4^. This adaptability ^5,6^ stems in part from its robust cell envelope ^7,8^, its ability to form biofilms ^9^, and its efficient exchange of genetic material via horizontal gene transfer (HGT), facilitating exceptional genomic plasticity ^10,11^. HGT is a key driver of *A. baumannii* genome evolution ^11–15^, enabling the rapid acquisition and dissemination of traits that promote survival ^17^, adaptation ^1,11,16–18^, and virulence ^19^. Understanding the mechanisms that underlie genomic plasticity and HGT is therefore essential to unravel the evolutionary strategies that contribute to the resilience of this pathogen.

The three canonical mechanisms of HGT in bacteria are conjugation, natural transformation, and transduction ^12^. In Gram-negative bacteria, conjugative transfer is facilitated by multicomponent type IV secretion systems ^20,21^, which typically assemble thin, flexible conjugative pili that promote donor–recipient cell contact and mating-pair formation ^21^. Conjugative plasmids, such as the *Escherichia coli* F-plasmid, can exceed 100 kb in size and be efficiently transferred between cells ^22^. In high-frequency recombination (Hfr) strains, in which a conjugative element is integrated into the bacterial chromosome, the conjugative machinery can mobilize large chromosomal segments in a single transfer event. This phenomenon has been extensively characterized in *E. coli* and demonstrated in other species such as *Salmonella* spp.^23^. However, to our knowledge, Hfr-mediated chromosomal transfer has not been demonstrated in *A. baumannii*. In natural transformation, bacteria take up environmental DNA and can integrate it into their genomes by homologous recombination. Natural transformation has been experimentally shown in several *A. baumannii* strains ^24–26^, with transformation frequency varying substantially between strains ^27^ and recombination tracts ranging from 13 to 123 kb based on genome-wide analyses of transformants ^24,28,29^. The third canonical mechanism of horizontal DNA transfer is transduction, where bacterial DNA is injected into a recipient bacterium via viral transducing particles ^30^. Generalized transducing phages often carry bacterial DNA fragments on the order of ∼5–100 kb; the size of DNA that can be transduced is typically constrained by phage capsid capacity, which depends on the phage family and capsid architecture ^31^. *A. baumannii* genomes contain a diverse repertoire of prophages encoding numerous antibiotic-resistance genes, and experimental studies have demonstrated prophage-mediated transfer of chromosomal resistance determinants between strains ^32,33^.

More recently, several non-canonical types of DNA transfer between bacteria have been described. These include outer membrane vesicle-mediated transfer, phage-like gene transfer agents, and membrane nanotubes ^12^. Outer membrane vesicles released by Gram-negative bacteria can encapsulate plasmid or chromosomal DNA fragments, typically ranging in the tens of kb ^34,35^. Gene transfer agents are phage-like particles that package random chromosomal fragments and generally transfer short DNA segments, typically a few kilobases in size ^36,37^. Membrane nanotubes have been observed in multiple bacterial species, including Gram-positive bacteria such as *Bacillus subtilis* ^38^ and Gram-negative species such as *E. coli* ^39^. In Gram-negative bacteria, these structures arise from extensions of the outer membrane and form lipid-based hollow connections between neighboring cells, which often appear as chains of vesicle-like structures with diameters of 40–80 nm and lengths up to tens of micrometers. Nanotubes are associated with the exchange of cytoplasmic molecules such as metabolites and proteins ^39–41^. Although transfer of non-conjugative plasmids between nanotube-connected cells has been reported ^38^, the size of DNA transported has not been determined and the passage of DNA through nanotubes has not been directly visualized. In addition, cell-cell bridges have been reported in the halophilic archaeon *Haloferax volcanii* ^42^. Unlike membrane nanotubes, which derive from membrane protrusions, these bridges are bounded by a continuous cell envelope and establish direct cytoplasmic continuity between neighboring cells. In *H. volcanii*, such bridges have been observed during cell–cell interactions and may facilitate exchange of cytoplasmic components, although DNA transfer through these bridges has not been demonstrated. Their existence suggests that direct cytoplasmic connections could provide an additional route for intercellular material transfer beyond classical bacterial HGT mechanisms. However, the functional significance of these bridges is poorly understood.

Here, we report the serendipitous discovery of intercellular membrane connections in *A. baumannii* that resemble conduits. We used cryo-electron tomography, super-resolution microscopy, and genetic and genomic analyses to investigate intercellular conduits found in clinical isolates of *A. baumannii*. Our analyses demonstrate that these conduits establish direct cytoplasmic continuity between neighboring cells and contain chromosomal DNA being transferred. Interstrain mating assays coupled with whole-genome sequencing revealed the transfer and homologous recombination of chromosomal segments up to 1.1 Mbp in length. We demonstrate that this transfer is distinct from natural transformation and conjugation. Together, these findings identify a conduit-mediated route for large-scale chromosomal DNA exchange with the potential to form chimeric pathogens with novel combinations of both virulence determinants and antimicrobial resistance factors.

## Results

### *A. baumannii* clinical isolates form intercellular membrane connections

We used electron microscopy to investigate the cell envelope of *A. baumannii* and visualize its protein constituents. During this analysis, we observed intercellular membrane connections. To investigate the occurrence and potential role of intercellular membrane connections in *Acinetobacter baumannii*, we analyzed the bacterial envelopes of the laboratory strain ATCC 17978 and its derivatives M317 and SC2151 and compared them with the envelopes of multidrug-resistant clinical isolates belonging to prevalent international clonal lineages (IC), including SC2072 (IC1), SC1842 (IC3), SC1833 (IC5), SC2076 (IC6), 1372 (IC2), 2778 (IC2), SC2073 (IC7), SC2075 (IC7), and SC1846 (IC3). All strains were cultured under identical growth conditions, fluorescently labelled with the lipophilic dye Vybrant DiO, and imaged using confocal fluorescence microscopy, with at least 400 randomly selected cells per strain. In some strains, the cells remained detached from one another (1372, 2778, SC2073, SC1846, and SC2075), but in others up to 7% of cells exhibited intercellular membranous connections of different lengths (Fig. 1a).

**Figure 1:**
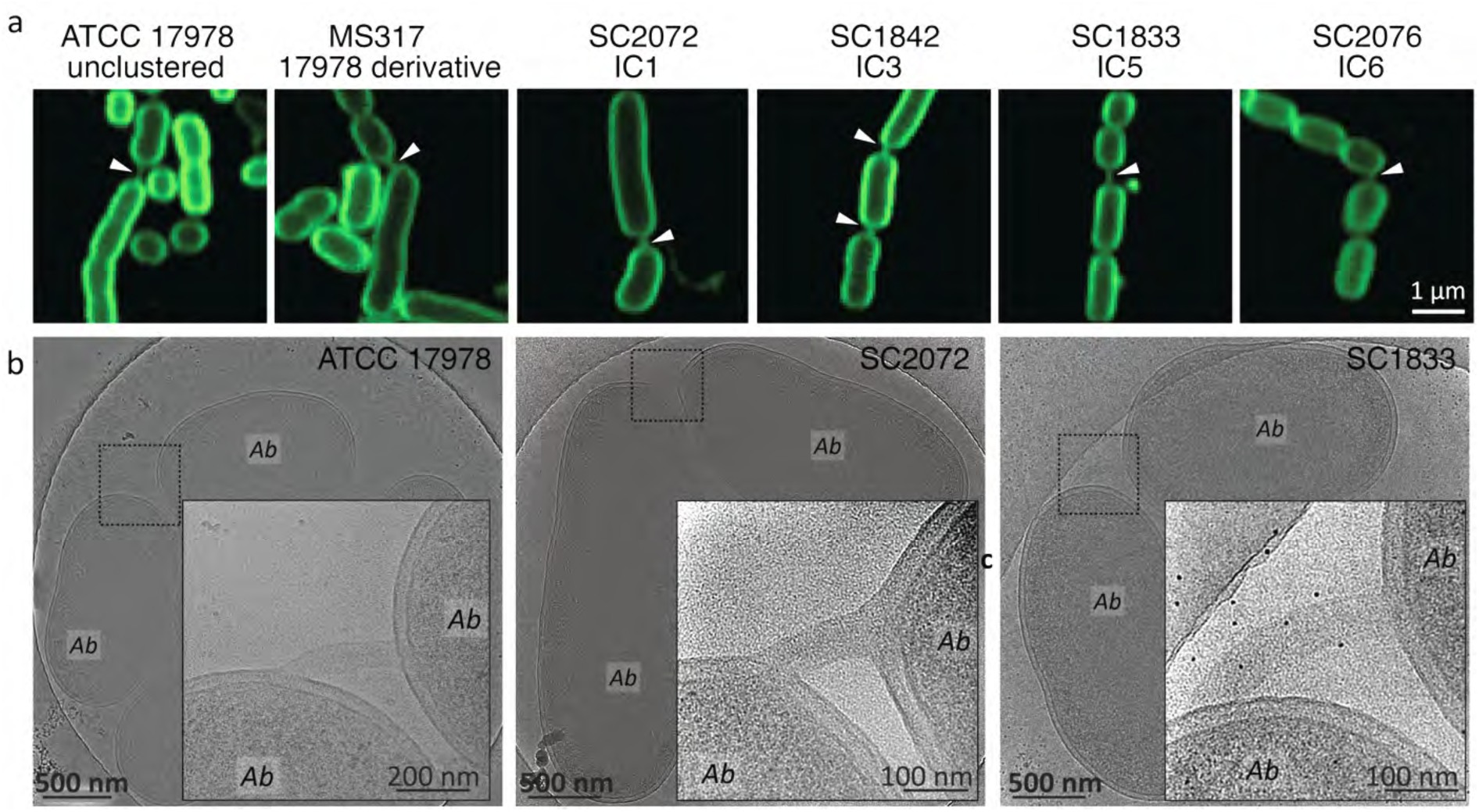
*A. baumannii* ATCC 17G78 and clinical isolates form membrane connections. **(a) Fluorescence light microscopy of six *A. baumannii* strains reveals the formation of membrane connections.** Cell membranes were labelled with the lipophilic dye Vybrant DiO. White arrows indicate the membrane connections. **(b) Cryo-electron micrographs show the membrane connections.** In the inset, the position of the membrane connection is magnified. Individual *A. baumannii* cells are labelled Ab.

We used cryo-electron microscopy (cryo-EM) of frozen-hydrated *A. baumannii* ATCC 17978 cells to characterize the observed membrane connections in greater detail. Consistent with our confocal fluorescence microscopy observations, we identified intercellular tubular membranous connections between the poles of *A. baumannii* cells (Fig. 1b). These connections were observed either between pairs of cells or between multiple cells arranged in a row (Fig. S1a, b). We also observed similar connections in the apathogenic species *Acinetobacter baylyi* from the same genus (Fig. S1c). Importantly, we could distinguish between division septa and intercellular membranous connections (Fig. S2). Notably, the connections in both species morphologically resemble the cell-cell bridges that have been, to our knowledge, thus far described only in Archaea ^44^.

We next quantified the occurrence of membrane connections in ATCC 17978 across growth phases (Table S1). For each phase, 250 randomly selected cells were analyzed, revealing an average frequency of 3.8% of cells displaying intercellular connections. Specifically, 3.3% of cells exhibited connections during exponential phase, 3.7% during early logarithmic phase, and 5% during late logarithmic phase. The connections were found at a substantially lower frequency (an average of ∼0.7% of cells during early logarithmic phase) in *A. baylyi* (2890 randomly selected cells, Fig. S1c). Notably, when cells were harvested directly from agar plates rather than liquid culture, the frequency of cells displaying intercellular membrane connections increased to as much as 19%. This indicates that growth conditions can substantially influence the formation of membranous connections (see Table S1 for detailed information), consistent with previous observations of growth condition-dependent formation of bacterial nanotubes ^43^.

### Membrane conduits in *A. baumannii* are continuous extensions of the cell envelope and contain filamentous structures

For cryo-electron tomography (cryo-ET) analyses, we used both native and mildly lysed (“ghosted”) *A. baumannii* cells to reduce the thickness and improve the resolution of the tomograms (see Materials and Methods). Mild lysis before cryo-ET was necessary, because (i) only a small fraction of cells formed membrane connections; and (ii) despite significant efforts, the membrane connections could not be found in the scanning electron microscope for focused ion beam milling, due to the poor resolution of the *in situ* light microscope.

All observed membrane connections were pleomorphic in curvature and length but had similar outer diameters (averaging 63.0 nm, σ = 5.5 nm, n = 95) (Fig. 2) and consistently originated from the pole regions of the *A. baumannii* cells. Cryo-electron tomograms revealed that these membrane connections appear as conduits formed by elongation of the cell envelope. The bacterial outer membrane, peptidoglycan layer, and inner membrane were clearly resolved within the conduits (Fig. 2), consistent with the canonical architecture of the Gram-negative envelope. The resulting inner lumen measured ∼10 nm in diameter (Table S1; Fig. 2). Examination of the base of the conduit, where it emerges from the cell envelope, did not reveal discrete large macromolecular complexes or obvious proteinaceous assemblies. Instead, the conduits formed continuous cytoplasmic bridges between adjacent cells (Fig. 2a, b). The conduits contained cytoplasmic material, including high-molecular weight complexes. In addition, we observed filaments running inside the conduit along its entire lumen (Fig. 2a, c, e). These filaments could be traced for >100 nm in length and were ∼2 nm thick, which is consistent with the diameter of double-stranded DNA (Fig. 2a, inset).

**Figure 2:**
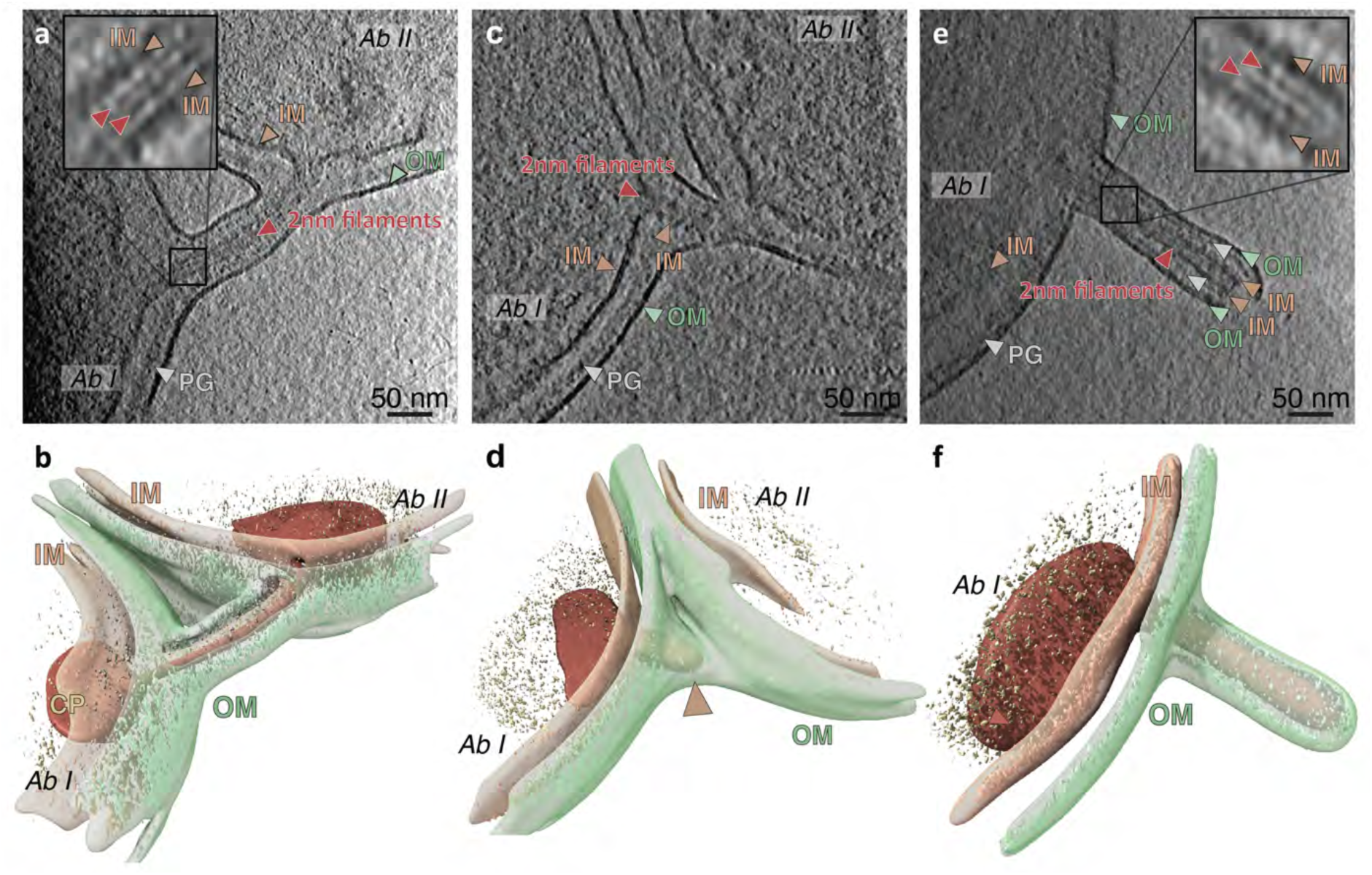
Cell envelope transformation associated with conduit formation revealed by cryo-electron tomography. (a,c,e) Tomographic slices (1 nm thickness) showing the outer membrane (OM, including the bilayer, green arrows), the peptidoglycan layer (PG, grey arrows), the inner membrane (IM, apricot arrows), the cytoplasm (CP, yellow arrows), and ∼2 nm thick filaments (brown arrows with a white boundary). Prior to plunge freezing, the cells were mildly lysed to reduce their thickness and improve resolution in the cryo-electron tomograms. (b,d,f) Isosurfaces and segmentation of bacterial components: OM (green), IM (apricot), CP (yellow), and PG (grey). The ∼2 nm thick filaments are indicated by brown spheres. (a,b) A membrane conduit, seamlessly formed by the IMs and OMs of two *A. baumannii* cells, connects their cytoplasm. The PG layer is continuous between the two cells. The conduit contains two ∼2nm-thick filaments that originate from the cytoplasm of the two connected cells. (c,d) A cell envelope extension originating from one *A. baumannii* cell, in which the OMs are fused while the inner membranes (IMs) remain unfused. The 2 nm-filaments can also be seen. (e,f) A ∼200 nm long extension of the cell envelope forming at the polar region of a single *A. baumannii* cell. Both inner and outer membranes are continuous with the cell envelope forming the extension. Within the extension, two ∼2nm-thick filaments can be seen. (a,e) The insets show projection slices of the filaments following sub-tomogram averaging, revealing the presence of two individual strands with a thickness of 2nm each, consistent with double-stranded DNA. The IM is indicated with apricot arrows, the two DA strands with brown arrows.

Similar filaments were detected within membrane conduits where only the outer membrane was fused to a partner cell (Fig. 2c, d) and in membrane extensions that had not yet established contact with neighboring cells (Fig. 2e, f). Thus, we hypothesize that these membrane extensions represent precursors that are not yet fully fused to a partner cell but that may later develop into intercellular conduits. In addition, the IM appeared decorated with electron-dense macromolecules, which were more readily apparent in the sub-tomographic projection slices (Fig. 2a, e, insets). The nature and molecular identity of these densities remain unclear.

### Membrane conduits contain DNA and display asymmetric nucleoid organization

To determine whether the membrane conduits indeed contain DNA, we combined stimulated emission depletion (STED) microscopy ^44^ with point accumulation for imaging in nanoscale topography (PAINT) ^45,46^ to image *A. baumannii* cells. Bacterial cells cultured for 4 h (early logarithmic phase) were immobilized on poly-L-lysine–coated chamber slides, membranes were labelled with Nile Red, and DNA was stained using JF_646_-Hoechst, as described previously ^46^. Super-resolution imaging revealed clear DNA signals overlapping with the membrane signal of the conduits (Fig. 3 a–b). In addition, DNA signals overlapped with membrane extensions at the cell poles that were not yet connected to a partner cell (Fig. 3c, d). Together, these observations indicate that the membrane conduits contain DNA.

**Figure 3:**
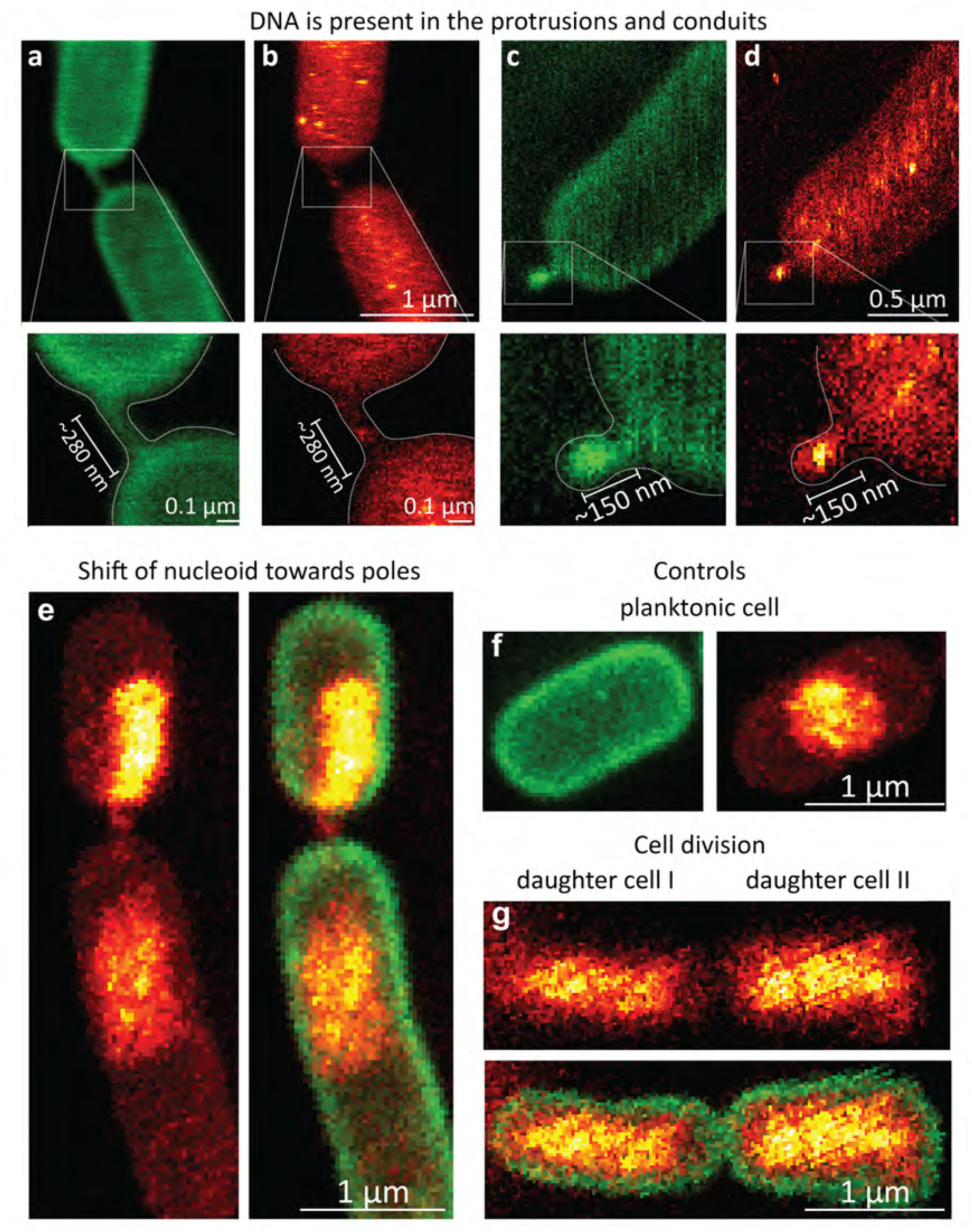
Super-resolution STED-PAINT microscopy reveals DNA within *A. baumannii* (ATCC 17978) membrane extension and intercellular conduits, with nucleoid displacement toward the conduit in one connected cell. Membrane staining was performed using Nile Red (shown in green) and DNA staining with JF_646_– Hoechst (shown as a red–yellow colormap, with bright yellow indicating higher signal intensity and dark red indicating lower intensity). Insets show magnified view. (a) Membrane conduit connecting two *A. baumannii* cells. The dimensions measured by STED-PAINT correspond to those of the conjugative bridges observed in cryo-electron tomography. (b) DNA signal detected within the conduit. (c) Membrane extension at the cell pole. (d) DNA signal detected within the membrane extension. (e) *A. baumannii* cells connected by a membrane conduit display the shift of one nucleoid toward the cell poles. This asymmetric nucleoid organization was characteristic for *A. baumannii* cells connected by membrane conduits. (f) Control experiment showing a Planktonic cell with a characteristic, centrally positioned nucleoid. (g) Control experiment showing a typical dividing *A. baumannii* cells, where the nucleoids are evenly distributed within the cell bodies.

We further observed highly asymmetric nucleoid localization in two cells connected by a conduit, with the nucleoid of one cell shifted toward the pole region (on average ∼36% of the total cell length). This pattern is consistent with the presence of distinct DNA-donating and DNA-receiving cells (Fig. 3e). This is further supported by images of one cell in a conduit-connected pair showing a centrally localized, typical nucleoid, while the nucleoid of the other cell is asymmetrically shifted towards the pole (Fig. S3). This asymmetric nucleoid localization contrasts with the centrally positioned nucleoid in planktonic cells (Fig. 3f), and the symmetric DNA distribution between daughter cells during division (Fig. 3g). Lastly, the existence of membrane protrusions that had not yet connected to partner cells is inconsistent with the conduits being remnants of failed cell division. Notably, the polar localization of the nucleoid and its association with the conduits prompt us to hypothesize that this mechanism may be specific to the transfer of genomic DNA.

### Conduit-mediated DNA transfer facilitates interstrain recombination and horizontal gene transfer

Next, we investigated whether the presence of DNA in the conduits and the polar localization of one nucleoid towards the conduit indicate a novel route for exchanging genomic information between bacterial cells. To test this, we co-cultured the conduit-forming *A. baumannii* strain SC2151 (95 of 1184 cells analyzed by cryo-EM formed conduits) and the non-conduit forming strain SC1846 (0 of 1479 cells analyzed by cryo-EM formed conduits, Fig. S4, Table S1), which are each resistant to a different antimicrobial agent (Fig. 4a). The *A. baumannii* SC1846 chromosome exclusively harbors the antibiotic resistance gene *bla*_OXA-23_, which confers resistance to meropenem. The chromosomes of both strains encode the preprotein translocase subunit SecA, albeit in different variants. The SecA variant in SC2151, which differs from the SC1846 variant by two amino acids, conveys resistance to sodium azide (NaN_3_) ^47^. After co-culturing, we identified, sequenced, and annotated a total of 22 bacterial clones that displayed a resistance profile distinct from either parental strain but combining the resistance phenotypes of both (Table S2), consistent with recombination.

**Figure 4:**
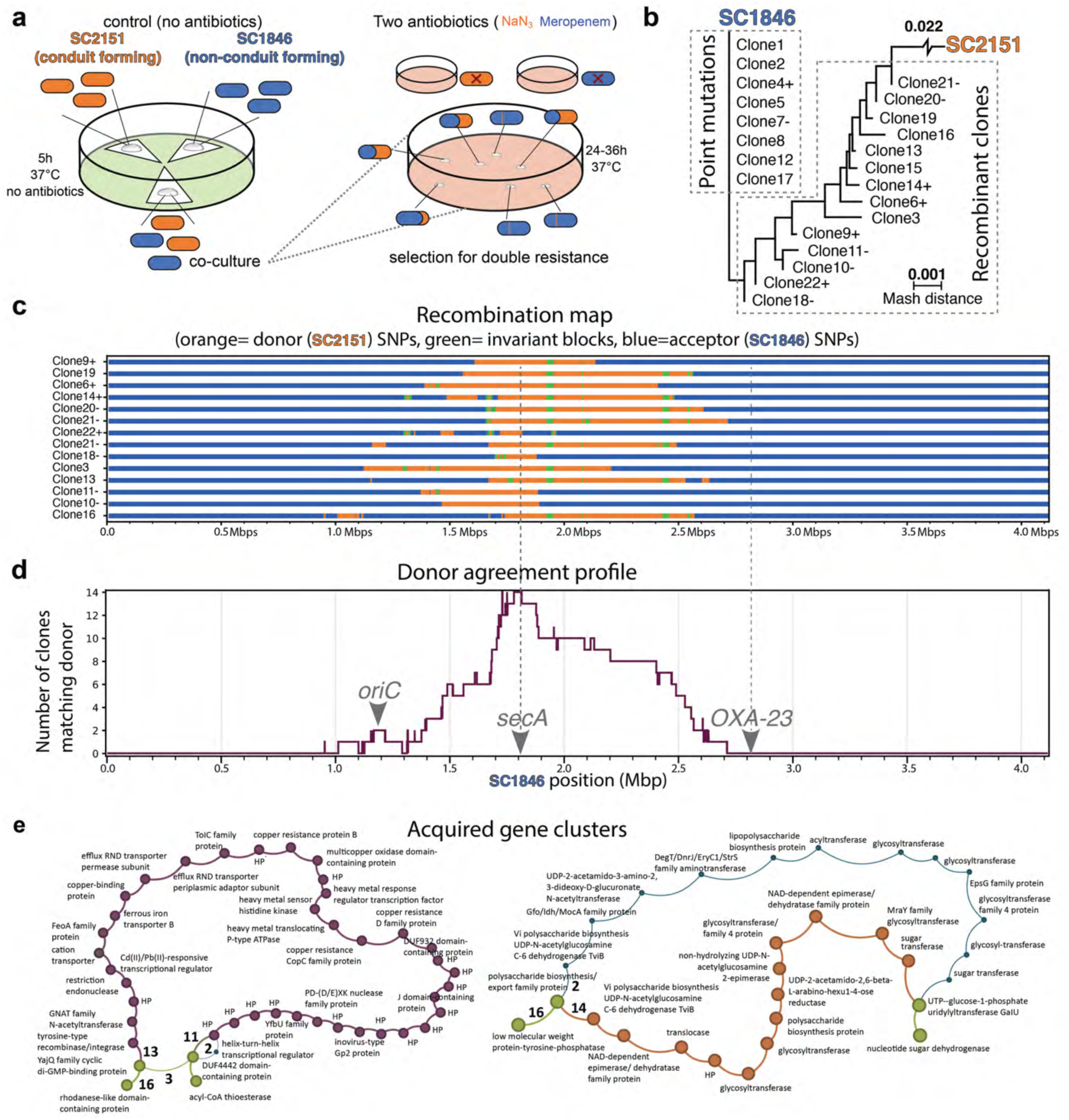
Conduit-mediated horizontal gene transfer, genomic recombination, and acquisition of functional traits. (a) Schematic representation of the bacterial mating experiment between the non-conduit forming strain SC1846 and the conduit-forming strain SC2151 during the HGT assay. (b) Hierarchical clustering of the parental strains (SC1846, SC2151) and the recombinant clones obtained from the HGT experiments using pair-wise Mash distances (genetic dissimilarity measure). Branch lengths approximate the genomic distance between pairs of taxa. 8 clones acquired genomic resistance due to point mutations (see also Fig. S5 and S6). The remaining 14 clones are recombinants, albeit closer to SC1846 than to SC2151. (c) Genome-wide recombination maps for the 14 recombinant clones using the SC1846 genome as reference. Variable positions reflecting the SC2151 genotype are indicated in orange and intervening invariant positions are colored in green (see also Fig. S7 and S8). Large contiguous blocks of sequence originating from SC2151 are integrated into the SC1846 genomic background, indicating extensive homologous recombination involving up to 1.1 million base pairs. (d) Cumulative plot summarizing the recombination events across all 14 clones shown in (c). Donor agreement profile derived from the recombination analysis shown in (c). The cumulative plot reaches the maximum at the locus encoding the NaN_3_ resistance-conveying SecA variant of SC2151. (e) Pan-genome graphs integrating gene content and order across 14 recombinant clones and the parental strains reveal HGT of functional modules. Nodes represent genes; node diameter indicates the number of genomes harbouring each gene. Edges connect genes that are adjacent in at least one of the 16 genomes, with edge weight indicating the number of genomes containing the gene pair. Edge weights are shown once per color and apply to all edges of that color. Green: core gene pairs present in all 16 genomes. Left: 12 recombinant clones acquired a heavy-metal-resistance gene cluster from SC2151 (violet nodes). Right: Exchange of a glycosylation-associated functional module. Orange nodes: SC1846 cluster; blue nodes: alternative SC2151 cluster present in one recombinant clone (Clone 10).

This analysis revealed two categories of clones. The first category comprised eight clones that are genetically almost identical to SC1846 (Fig. 4b). A dot plot showed that each of the eight clones was collinear to SC1846 (Fig. S5), and a multiple whole-genome sequence alignment revealed only 142 single nucleotide variants across all eight clones when compared to SC1846. Analysis of the *secA* locus revealed missense mutations in six of the eight clones. Importantly, these mutations differed from the *secA* variants present in the NaN_3_-resistant SC2151 (Fig. S6). Thus, the double-resistance phenotype in these clones likely arose from independent spontaneous point mutations within the SC1846 background and cannot be explained by a conduit-mediated DNA transfer. The second category comprised the remaining 14 clones, who were genetic intermediates between SC1846 and SC2151, albeit more similar to SC1846 than to SC2151 (Fig. 4b). In line with this latter finding, dot plots reveal locally confined regions of similarity to SC2151 within a genomic background resembling SC1846 (Fig. 4c, Fig. S7), and whole-genome sequence alignment identified a total of 31,669 and 74,563 single nucleotide variants when compared to SC1846 and SC2151, respectively. Taken together, these findings suggest that the clones in the second category are recombinants where genomic fragments from SC2151 were transferred to SC1846 and subsequently integrated into its genome.

To trace recombination events in the genomes of these 14 clones, we determined sites where the genotype switched from the variant seen in SC1846 to that of SC2151 and displayed them along the genomic sequence of SC1846 (Fig. 4c). Reproducing the findings from our initial analyses (see Figs. 4b, c) the clones resembled the genotype of SC1846, across most of their genomes. Importantly, the regions where the genotype of the clones more closely resembled that of SC2151 were not randomly distributed along the bacterial chromosome. Instead, we detected contiguous stretches of up to 1.1 million base pairs (200–1,100 kb). This is consistent with horizontal gene transfer of very large genomic segments from SC2151 to SC1846, followed by their recombination into the SC1846 genome (Fig. 4c, d). In all 14 clones, the *secA* locus was included in the recombined regions, and multiple-sequence alignment confirmed that all clones carried the SC2151 variant of this gene (Fig. S8).

The conduit-mediated recombination of more than a quarter of a bacterial chromosome has three possible outcomes for the genetic repertoire in these clones: for genes located in shared syntenic regions in the parental strains, it converts the acceptor variant to the donor variant (see Fig. S8 for an example). For genes that are exclusively present in one of the parental strains, the recombination event changes the gene set composition. It can either introduce donor genes into the genomic context of the acceptor strain (gain of function), or it can induce the deletion of genes that are originally present only in the acceptor (loss of function). To investigate the effects at the gene set level, we generated a pan-genome graph for the 14 recombinant clones and the two parental strains with PPanGGOLiN ^48^. In the individual recombinant clones, between 11 and 130 SC1846 gene families were replaced by 12 to 141 SC2151 gene families (Table S3). The net change in gene family content was small, with a mean gain of 21 families (minimum: -14; maximum: 45). This likely reflects the fact that the SC1846 and SC2151 chromosomes are largely co-linear (Fig. S7) and share most (70%) of their genes. Still, the functional impact of the local gene set turnover can be substantial. As an example of a loss of function, two adjacent SC1846 genes involved in aspartate metabolism are absent from two recombinant clones because the corresponding region in SC2151 contains different genes (Fig. S9). Conversely, this recombination event introduced SC2151 genes into the corresponding genomic interval, resulting in a gain of function. As a further example for a gain of function,12 of the 14 recombinant clones acquired a cluster comprising 38 genes, many of which are involved in heavy metal resistance (Fig. 4e, left). In a more complex scenario gene gain and loss at a locus can also be combined as it is demonstrated by the exchange of a “gene cassettes” comprising alternative pathways for glycosylation (Fig. 4e, right).

### Large-scale chromosomal transfer is distinct from natural transformation and pilus-dependent conjugative transfer

Distinguishing between natural transformation, conjugation, and conduit-mediated transfer is challenging, as the underlying mechanisms and the proteins mediating the observed DNA transfer remain unknown.

We performed co-culture experiments in the presence or absence of DNase, which yielded identical outcomes, arguing against uptake of free extracellular DNA as the transfer mechanism. Analysis of the plasmid content of strains SC2151 and SC1846 and 22 bacterial clones also revealed no evidence of additional plasmid transfer between the strains (Table S4).

To further assess the contribution of natural transformation, each strain was exposed to purified genomic DNA from the other strain. Double-resistant colonies were obtained from SC1846 exposed to SC2151 DNA, and 24 independent control isolates were analyzed. Whole-genome sequence alignment and single nucleotide variant analysis identified only 16 substitutional variants among the control isolates. None of these variants indicated the incorporation of donor-derived DNA or large-scale chromosomal transfer. Notably, all 24 control isolates shared the same nonsynonymous mutation in *secA*(E→K), which is associated with NaN_3_ resistance. However, this mutation differed from the corresponding allele in SC2151, indicating that it was not acquired from the donor strain. These results suggest that uptake of free extracellular DNA cannot explain the extensive DNA transfer observed in the co-culture experiments.

Genome analysis using OriTfinder did not identify a complete set of predicted conjugation genes in either strain. SC2151 encodes a predicted Type IV secretion system (T4SS) and T4 coupling protein (T4CP) but lacks a predicted oriT and relaxase, whereas SC1846 encodes a relaxase and T4CP but lacks a predicted oriT and T4SS. Thus, neither strain appears to encode a complete canonical conjugation system, arguing against classical T4SS-mediated, pilus-dependent conjugative transfer as the mechanism underlying the observed DNA exchange (Fig. S10).

To assess whether conduit formation could be attributed to a specific genetic determinant, we compared the genomes of conduit-forming *A. baumannii* strains SC2151, ATCC 17978, SC1833, and SC2072, as well as *A. baylyi*, with that of the non-conduit-forming strain SC1846. This comparison did not identify any individual gene or gene cluster that was consistently associated with the conduit-forming phenotype.

## Discussion

*Acinetobacter baumannii* exhibits extraordinary genomic plasticity, enabling rapid adaptation to diverse and often hostile environments, including hospitals ^1,2,9^. Here we describe intercellular conduits that establish direct cytoplasmic connections between neighboring cells and mediate extensive genome exchange. Through this pathway, chromosomal DNA segments of millions of base pairs can be transferred between clinical strains, thereby revealing a scale of genetic plasticity far beyond canonical horizontal gene transfer mechanisms in *A. baumannii* ^22,24,31,34,36^.

The conduits described here represent extensions of the entire cellular envelope and differ fundamentally from previously described extracellular structures. In contrast to outer membrane vesicles, bacterial nanotubes ^49^, or other projections that bud from the outer membrane and consist of a single membrane layer ^50^, these conduits consist of both membranes and the peptidoglycan layer. They are also distinct from proteinaceous appendages such as pili ^51^. Bacterial nanotube formation has been associated with a conserved membrane-associated complex that shares components with the flagellar basal body ^38,43,52^. However, *A. baumannii* lacks flagellar genes ^53^, making involvement of this flagellar-associated machinery in conduit formation unlikely in this organism. Morphologically similar conduits have been observed in Archaea ^42^, but DNA has not been seen crossing them.

Several observations argue against the possibility that conduits represent remnants of incomplete cell division. Conduit-connected cells consistently displayed asymmetric nucleoid positioning toward the conduit, and we frequently observed isolated membrane protrusions that had not yet established contact with another cell. In addition, conduit frequency did not differ significantly across the examined growth phases, although it increased markedly when cells were harvested from agar plates.

The two filaments of double-stranded DNA observed within the conduits and in the protruding conduit precursors suggest that genetic material is priorly attached to the inner membrane during conduit formation. Our genomic analyses provide independent evidence for large-scale DNA exchange, with homologous recombination of large chromosomal segments spanning more than a million base pairs and generating mosaic genomes that incorporate contiguous donor-derived regions containing resistance-associated loci. Our analysis of recombination boundaries suggests that DNA transfer and/or recombination may preferentially initiate at specific genomic locations. In most recombinants, the transferred fragments terminate immediately downstream of *secA*, the locus used for selection. The respective other terminus can be as far as 1 Mbp apart from the SecA locus, with several instances clustering at around 2.5 Mbp (see Fig. 4c, d). Notably, all detectable recombination events originated from strain SC2151 and were incorporated into strain SC1846. This directionality is consistent with our observation that SC2151 forms membrane conduits at a frequency of ∼6%, whereas no conduits were detected in SC1846, suggesting that SC2151 may serve as the DNA donor and SC1846 as the recipient in conduit-mediated transfer. However, because the mating experiments relied on selection for both the SC2151 *secA* allele and the SC1846 *bla*OXA-23 gene, we cannot exclude the possibility that transfer events in the opposite direction would remain undetected under the present experimental design. This could occur if the putative origin of transfer or recombination lies upstream of the blaOXA-23 gene.

Importantly, DNase treatment did not reduce transfer, and exposure to purified genomic DNA produced only small numbers of control recombinants lacking donor-derived large chromosomal tracts, arguing against natural transformation as the mechanism underlying the observed large-scale transfer ^24–26^. Likewise, the structural and functional characteristics described here are not readily explained by canonical pili-based conjugation ^54,55^. In classical type IV secretion system-mediated conjugation, DNA is typically transferred as a single-stranded molecule through a dedicated secretion apparatus. By contrast, the conduits described here form continuous membrane-bound cytoplasmic bridges containing genomic DNA, as supported by STED-PAINT, asymmetric nucleoid positioning, and transfer of large chromosomal segments; the ∼2-nm-thick filaments within the conduits are consistent with double-stranded DNA. While neither strain appears to encode a complete canonical conjugation system, we cannot exclude the possibility that individual elements of the conjugation machinery contribute to conduit formation or stabilization.

The molecular machinery underlying conduit biogenesis remains unknown. Examination of the conduit bases in the cryo-EM images did not reveal large macromolecular complexes or obvious protein assemblies that might drive conduit formation. Electron-dense macromolecules were observed along the inner membrane, but their identity remains unclear. Comparative genomic analysis also did not identify any gene or gene cluster consistently associated with the conduit-forming phenotype. Thus, conduit formation may involve smaller, transient, or strain-specific factors that are not readily detectable by our approaches.

Together, our findings identify conduit-mediated chromosomal gene exchange as a mechanism of horizontal gene transfer in *A. baumannii*. By enabling the transfer of large genomic regions between clinical lineages, this process provides a potential route for the dissemination of antimicrobial resistance and may contribute to the remarkable evolutionary adaptability of this opportunistic pathogen.

## Acknowledgements and Funding

We thank Daniela Bublak and Sara Riedel-Christ for the cultivation of *A. baumannii*. We thank the Frankfurt Center for Electron Microscopy and the Frankfurt Center for Advanced Light Microscopy for measurement time. C.B. was funded by Bundesministerium für Bildung und Forschung (BMBF, Germany) in partnership with the l’Agence Nationale de la Recherche (ANR, France) (program EFFORT, ANR-19-AMRB-0007, BMBF-16GW0236K/01KI2123 (K.M.P., A.S.F.); A.S.F. was supported by the Deutsche Forschungsgemeinschaft (Research Training Group iMOL GRK 2566, project number 414985841, for S.M. and P.R. and FR1653/6-3); A.B. and M.H. were supported by the Deutsche Forschungsgemeinschaft (Research Training Group iMOL GRK 2566, project number 414985841; CRC 1177, project number 259130777; INST 161/1020-1 FUGG); J.S. was funded by the Deutsche Forschungsgemeinschaft (project no. 493624332); F.L. and I.E. were supported by the Marie Curie ITN project StraDiVarious (Project-ID: 101226609); F.L. and I.E. were supported by the Marie Curie ITN project StraDiVarious (Project-ID: 101226609); S.G: was supported by the Dr. Rolf M. Schwiete-Stiftung; MH, IH, KMP, ASF acknowledge funding by the Deutsche Forschungsgemeinschaft (DFG, German Research Foundation) under Germanýs Excellence Strategy – EXC-3094 – 533751785.

## Author Contributions

S.M.: Designed experiments, planned and carried out *A. baumannii* mating experiments, performed HGT assays. C.B.: Designed experiments, planned and carried out *A. baumannii* sample preparation, recorded and processed cryo-EM and cryo-ET datasets, and performed light microscopy experiments. S.M. and P.R.: Carried out *A. baumannii* sample preparation, recorded and analyzed cryo-EM datasets. C.B. and A.B.: Carried out super-resolution microscopy experiments. M.H.: Supervised super-resolution microscopy experiments. S.G.T. and C.T.: Performed HGT assays and genome sequencing. S.G.: Selected and provided clinical *A. baumannii* isolates, carried out genomic sequencing and gene analysis, and supervised research. I.E. designed bioinformatics analyses and supervised research; J.S., F.L., P.R. and I.E.: Performed genome sequence analysis and comparative gene set analyses. A.S.F., I.H. and K.M.P.: Designed and planned experiments, and supervised research. S.M., C.B. and A.S.F.: Wrote the manuscript, with contributions from all authors.

## Data Availability

Whole-genome sequences of the conduit-forming and non-conduit-forming strains generated in this study have been deposited in DDBJ/ENA/GenBank under BioProjects PRJNA1516054, PRJNA1328187, PRJNA1255660 and PRJNA901493. Individual BioSample accession numbers for all isolates are listed in Table S5. The genome of the reference strain *A. baumannii* ATCC 17978 is available under BioSample SAMN02604331.

## Materials and Methods

### Bacterial strain culturing

The *A. baumannii* strain American Type Culture Collection (ATCC) 17978 used in this study was kindly provided by Dr. Higgins (DZIF, Cologne). For control experiments, the reference strain ATCC 17978 and clinical isolates were cultured and prepared by the laboratory of Prof. Göttig (University Hospital Frankfurt). Bacterial cultures were grown by selecting bacterial colonies from LB agar plates and transferring them into Luria–Bertani (LB) medium. Bacterial pre-cultures were grown overnight at 37 °C. After 12 h, the main cultures were inoculated at an optical density (OD_600_) of 0.05-0.1 and grown at 37 °C until the desired OD_600_ was reached.

### Confocal laser scanning microscopy

A total of 2 mL of bacterial cells grown to the desired OD_600_ were harvested by centrifugation at 4500 *xg* for 3 min at room temperature (RT). The resulting pellets were resuspended in (phosphate-buffered saline (PBS) and inactivated with 6% electron microscopy (EM)-grade methanol-free paraformaldehyde (Electron Microscopy Sciences) at RT. The cells were then washed with PBS and labelled with the fluorescent membrane dye Vybrant DiO (Thermo Fisher Scientific) at a concentration of 1:60 at 30 °C for 30 min under light exclusion. After labeling, the cells were centrifuged (4500 *xg*, RT, 2 min) to remove excess dye and washed once with PBS. Cells were then resuspended in PBS and imaged on agarose gel pads (1% low-melt agarose in 89 mM Tris-borate, 2 mM EDTA, pH 8.0). Confocal fluorescence images were acquired using a laser scanning microscope (Zeiss LSM 700) with a Plan-Apochromat 63×1.40 Oil DIC objective and Zeiss LSM software. The LED laser (488 nm) was used at an intensity of 0.3%, and the signal was captured with a photomultiplier tube (PMT) detector using a SP555 filter. Gain settings were individually adjusted for each sample, ranging between 400 and 600 V. The pinhole was set to 1 airy unit (44.6 μm), and images were acquired with a pixel size of 0.09 µm and a pixel dwell time of 2.85 µs. To maximize the signal-to-noise ratio, averaging of two scans was performed line by line. The acquired images were processed using the Fiji software ^56^.

### Cryo-electron microscopy sample preparation of native *A. baumannii* cells

After the bacterial cells reached the desired OD_600_, 2 mL of culture were harvested by centrifugation at 4300 ×g for 3 min at RT. The pellets were resuspended in 500 µL of LB medium and inactivated with EM-grade methanol-free paraformaldehyde (Electron Microscopy Sciences) at a final concentration of 6% for 1 h at RT. The inactivated cells were then centrifuged (4300 ×g, RT, 3 min) and resuspended in 50–150 µL of LB medium. Alternatively, cells were collected directly from LB agar plates, resuspended in 100 µL PBS, and inactivated with 6% EM-grade methanol-free paraformaldehyde (Electron Microscopy Sciences) for 1 h at RT.

For cryo-electron microscopy, 3.5 µL of sample were deposited onto glow-discharged Quantifoil R3.5/1, 200-mesh Cu holey grids, with or without a carbon film, and vitrified by plunging into liquid ethane using a Vitrobot Mark IV (Thermo Scientific, Waltham, USA) at 100% relative humidity and 4 °C, with a nominal blot force of −2 and wait and blotting times of 8 s. Alternatively, grids were blotted only from the backside using a Teflon sheet on the front, at 100% relative humidity and 4 °C, with a nominal blot force of −2, a wait time of 5 s, and a blotting time of 60 s. Depending on the experimental approach, 5-nm gold beads conjugated to protein A (CMC, Utrecht) were added to the samples as fiducial markers. All pipetting steps were performed using a cut-off pipette tip to avoid damaging the membrane conduits.

### Cryo-electron microscopy sample preparation of *A. baumannii* ghost cells

After reaching the desired OD_600_, the bacterial cells were taken from the culture with a final concentration of OD_600_ = 5 and added to equal volumes of LB medium and cold H_2_O to reach a final volume of 10 mL. The cells were harvested for 15 min at 4500 *xg* and 4 °C. After three washes with 50 mM EDTA and 20 mM Tris buffer in a volume of 1 mL at 4300 *xg* for 2 min at RT, the cells were transferred to 1 mL of 20 mM Tris ice-cold buffer and washed five times at 4300 *xg* for 2 min at RT. Thereafter, the cells were resuspended in a mixture of 1 mL 1.5 M sucrose and 1.7 mL 0.9 M sucrose, and lysozyme was added to the sample to achieve a final concentration of 250 µg/mL. The suspension was incubated for 2 min on ice and 3 min at RT and inverted regularly. Then, 25 mL of ice-cold PBS (pH 7.4) was added to the cell suspension, and the suspension was gently stirred for 15 min in an ice bath. To inactivate the bacterial cells, EM-grade methanol-free paraformaldehyde (Electron Microscopy Sciences) was added to achieve a final concentration of 6% to 1 mL of the cell suspension on ice. Glow-discharged Quantifoil R3.5/1, 200-mesh Cu holey carbon-coated grids were placed in centrifugation inserts, and 83 µL of sample was applied to each grid. The cells were centrifuged onto the grids at 4500 *xg* for 45 min at 4 °C, with no acceleration and no deceleration. After centrifugation, the grids were washed in KCl buffer (pH 8) to remove any remaining sucrose. Finally, the grids were vitrified in liquid ethane using a Vitrobot Mark IV at 100% humidity, 4 °C, nominal blot force -3, a wait time of 0 s and a blotting time of 15 s. Depending on the experimental approach, 5 nm gold beads conjugated to protein-A were added to the sample as fiducial markers. All pipetting steps were performed using a cut-off pipette tip to avoid damaging the membrane conduits.

### UV-C inactivation of frozen cryo-EM grids

To exclude any potential effects of the paraformaldehyde fixation on the biological sample, UV-C inactivation of the *A. baumannii* cells was applied as an alternative treatment, following an UV-C inactivating approach similar to that described by Depelteau *et al*. ^57^ (IV44-53r30.03UFM122.21.02; 9.99.13/04 with data of 13.01.2022).

### Cryo-electron microscopy data collection

Micrographs were collected using SerialEM v4.2.0 beta ^58^ at a nominal magnification of 11,500× (16.4 Å per pixel) in nanoprobe mode on a Titan Krios G3i transmission electron microscope (Thermo Fisher Scientific) operated at 300 kV and equipped with a GIF Quantum S.E. post-column energy filter in zero loss peak mode and a K3 direct electron detector (Gatan). Each micrograph was recorded with a total dose of 1.5 e⁻/Å². The camera was operated in counting mode at a dose rate of 32 e⁻/px /s. Data were collected at a defocus of approximately −50 µm.

Tilt series were acquired with SerialEM v3.8 ^58^ on a Titan Krios transmission electron microscope (Thermo Scientific) operating at 300 kV. The nominal magnification was set to 81,000x (0.9 Å per pixel) in nanoprobe EFTEM mode. The microscope was equipped with a GIF Quantum S.E. post-column energy filter in zero loss peak mode and a K2 Summit detector (Gatan Inc., Pleasanton, USA). A total dose of 130 e-/Å^2^ per tomogram was applied. The tilt series covered an angular range from -60° to 60°, with an angular increment of 3° and a defocus set to -3 µm. The acquired tomograms were reconstructed by applying super-sampling SART ^59^. Electron micrographs for 2D visualization were recorded at a nominal magnification of 81,000x (0.9 Å per pixel) or 64,000x (1.1 Å per pixel). The total dose ranged from 30 to 45 e-per A^2^s^-1^, with a frame time of 0.2 s at a defocus varying from -3 µm to -5 µm.

### Whole-cell STED-PAINT microscopy

Sample preparation and stimulated emission depletion (STED) microscopy were performed as described previously ^46^. After *A. baumannii* reached an OD_600_ of 2.7, aliquots of the culture were harvested (6000 *xg*, RT, 2 min) and resuspended in LB medium. Cells were then fixed in solution for 1 h at RT by adding a mixture of 6% EM-grade methanol-free paraformaldehyde (Electron Microscopy Sciences) and 1.5% EM-grade glutaraldehyde (Electron Microscopy Sciences). Subsequently, the cells were centrifuged again (2 min, 6000 *xg*) and resuspended in PBS containing 0.2% sodium borohydride for quenching excess aldehydes for 3 min. The cells were then washed three times with PBS (6000 *xg*, RT, 2 min) and immobilized on 8-well chamber slides (Sarstedt) that were pre-treated with 3 M KOH (60 min at 40 °C) and coated with poly-L-lysine (Sigma). Additional post-fixation was performed using 2% EM-grade methanol-free paraformaldehyde for 10 min, followed by quenching of excess paraformaldehyde using 50 mM ammonium chloride in PBS for 10 min. After cells were washed three times with PBS, they were permeabilized using 0.5% Triton X-100 (Sigma) in PBS for 30 min and then rinsed three times with PBS. The *A. baumannii* membranes were labeled using Nile Red (Sigma, catalogue number N3013), and *A. baumannii* DNA was labeled using JF_646_-Hoechst. Stock solutions of Nile Red and JF_646_-Hoechst (Methanol for Nile Red and 100 µM in DMSO for Hoechst conjugate) were diluted in 150 mM Tris (pH 8.0). A final dye concentration of 300 nM was used for STED microscopy. STED microscopy was performed on a Abberior Expert line system (Abberior Instruments, Göttingen, Germany) operated by the Lightbox interface of the Imspector software (v16.3.15521-w2209; Abberior Instruments). Samples were imaged with an Olympus UPLXAPO60x NA 1.42 oil immersion objective (UPLXAPO60XW/1.42; Olympus, Japan) on an IX83 stand (Olympus). Nile Red was excited with a 560 nm pulsed laser (8%, 1.25 µW at the back focal plane) and JF_646_-Hoechst with a pulsed 640 nm laser (20%, 31.3 µW at the back focal plane). Stimulated emission depletion was performed with a 775 nm pulsed laser (Nile Red: 20%, 220 mW at the back focal plane; JF_646_-Hoechst: 25%, 270 mW at the back focal plane). Imaging was performed in a frame sequential mode for 2D-STED microscopy and in a line sequential mode for 3D-STED microscopy. The fluorescence signals for both channels were detected on two avalanche photo diodes (APDs) with a bandpass filter of 570–625 nm for Nile Red and 650–730 nm for JF_646_–Hoechst. A gating of 0.75 ns was applied. The pinhole was set to 0.81 AU and the pixel dwell time to 10 μs in the case of Nile Red and 5 µs in the case of JF_646_-Hoechst. Each line was scanned 70 times, and the signal was accumulated. Pixel size was set to 10 nm for 2D-STED microscopy and 20 nm for 3D-STED microscopy. We used a section depth of 80 nm between planes for 3D-STED microscopy. Image analysis was performed using the open-source image analysis package Fiji ^56^.

### HGT assay

Starter cultures of *A. baumannii* SC2151 and SC1846 were inoculated from single colonies into 10 mL of LB medium and incubated for 24 h at 37 °C. The medium contained 150 µg/mL sodium azide (NaN₃) for SC2151. On the following morning, the OD₆₀₀ of both cultures was measured and adjusted to an OD_600_ of 0.05 for SC1846 and of 0.1 for SC2151, and the cultures were incubated for 2 h at 37 °C. Cultures were then washed by centrifugation and resuspended in fresh LB medium to a final OD₆₀₀ of 2. For conjugation, three sterile filter paper pieces were placed onto a blood agar plate. Onto each filter, 12 µL of cell suspension was added as follows: (1) 12 µL of SC2151; (2) 12 µL of SC1846; or (3) a 2:1 mixture consisting of8 µL of SC2151 and 4 µL of SC1846. The plate was incubated upside down for 5 h at 37 °C to allow for conjugation. To control for natural transformation, 40 µg/mL DNase was added to the medium in some of the replicates.

After incubation, each filter was transferred into a sterile 1.5 mL tube containing 1 mL of sterile PBS and vortexed thoroughly to resuspend the cells. For the mixed (conjugation) sample, 100 µL of the undiluted suspension was plated, along with serial dilution at 10⁻³ on LB plates containing both 150 µg/mL NaN₃ and 2 µg/mL meropenem. For the SC2151 and SC1846 monocultures, 100 µL of the undiluted suspension and a 10⁻² dilution were plated as a negative control on the selective plates. To determine total colony-forming units (CFUs), serial dilutions of each strain (10⁻⁶, 10⁻⁹) were also plated on blood agar plates. Colony growth was monitored on the plates over the following days. In control plates containing both 150 µg/mL NaN₃ and 2 µg/mL meropenem, no colony growth was observed for either SC2151 or SC1846 alone. Colonies that emerged on the selective conjugation plates were picked, and their resistance profile was characterized by comparing inhibition zone diameters in a disk diffusion assay. Clones showing antibiotic susceptibility traits of both donor and recipient were subjected to further testing via colony PCRs to directly probe the presence of OXA-23. Colonies whose resistance profile matched both SC2151 and SC1846 were then subjected to whole-genome sequencing as described above.

### DNA extraction, whole-genome sequencing and bioinformatic methods

Whole-genome sequencing of all strains was carried out using short-read (MiSeq, Illumina) and/or long-read (MinION, GridION, PromethION, Oxford Nanopore) sequencing as described previously ^60^.

For the 22 recombinant and control isolates analyzed in this study, bacterial isolates were cultured on Columbia blood agar plates for 18 h at 37 °C. DNA was extracted using the DNeasy UltraClean Microbial Kit (Qiagen, Hilden, Germany) according to the manufacturer’s instructions. Whole-genome sequencing was performed using long-read sequencing on the PromethION platform (Oxford Nanopore Technologies, Oxford, UK). Library preparation was carried out using the SQK-NBD114.96 ligation sequencing kit (Oxford Nanopore Technologies). Sequencing was conducted on a P2 Solo PromethION, GridION or MinION instrument using R10.4.1 flow cells. Raw signal data was base-called and demultiplexed using the super-accuracy model of the Dorado basecaller v1.2.0. Raw data was trimmed using trimmomatic v0.39 for short reads and NanoFilt v2.8.0 for long reads. *De novo* assembly was conducted using Unicycler v0.4.8. and Autocycler v0.5.2 ^61^.

*Genome* annotation was carried out with PGAP v6.10 ^62^. Identification of antimicrobial resistance genes was performed using ABRicate v1.0.1 with the NCBI AMRFinderPlus database. Strain phylogeny was constructed using mashtree v1.4.6 ^63^, utilising Mash distance calculations, and visualised using the Phylo module from BioPython v1.86 ^64^. Whole genome alignments were performed with Mauve snapshot_2015-02-25 ^65^, substitutional variants were exported from the Mauve alignment and recombinant regions were identified and mapped onto the SC1846 reference genome with a custom python script. Pan-genome construction of annotated assemblies was performed using PPanggolin v2.2.6 ^48^, and gene presence-absence was calculated across all assemblies.

