## Supplemental Information for "Cell-envelope conduits enable transfer of megabase-sized double-stranded DNA between cells of the nosocomial pathogen *Acinetobacter baumannii*"

**Table S1: Membrane conduits quantified by growth phase and preparation conditions.** Conduits and envelope extensions were counted separately. Growth phases: Exponential phase (Exp.), early stationary phase (Early stat.), late stationary phase (Late stat.); Inactivation methods: paraformaldehyde fixation (PFA) or UV-C light (see Materials and Methods).

**(a)** Number of membrane conduits (MCs) observed in *A. baumannii* ATCC 17978. To exclude any potential effects of PFA fixation on MC formation, cells at the same growth stage ($OD_{600}$ = 0.5) were inactivated with either PFA or UV-C and screened independently, resulting in a similar total number (11 and 10, respectively) of observed MCs. Likewise, mild lysis did not alter the number of MCs (11) at the same growth stage ($OD_{600}$ = 2.5). Overnight cultures are indicated with (o/n).

**(b)** Number of MCs observed in *A. baumannii* ATCC 17978 from another laboratory (University Hospital Frankfurt) cultivated and prepared under identical conditions in triplicate.

**(c)** Number of MCs observed in *A. baumannii* ATCC 17978 harvested directly from culture plates.

**(d)** Number of MCs observed in *A. baylyi* ATCC 17978 (wild type; WT) harvested directly from culture plates.

**(e)** Number of MCs observed in *A. baumannii* SC2151 harvested directly from culture plates.

**(f)** Number of MCs observed in *A. baumannii* SC1846 harvested from liquid culture or directly from culture plates.

| | Strain | $OD_{600}$ | Growth phases | No. of counted cells per sample | No. of *A. baumannii* membrane connections | | | Prep. |
|---|---|---|---|---|---|---|---|---|
| | | | | | Conduit | Envelope extension | Total | Inacti-vation |
| **(a)** | *A.b.* ATTC 17978 | 0.5 | Exp. | 250 | 8 | 3 | 11 | PFA |
| | | 0.5 | Exp. | 250 | 7 | 3 | 10 | UV-C |
| | | 1 | Exp. | 250 | 9 | 0 | 9 | PFA |
| | | 1.7 | Exp. | 250 | 9 | 0 | 9 | PFA |
| | | 2.5 | Early stat. | 250 | 10 | 1 | 11 | PFA |
| | | 2.5 | Early stat. | 250 | 7 | 0 | 7 | PFA |
| | | 3.5 | Late stat. | 250 | 19 | 1 | 20 | PFA |
| | | 4 (o/n) | Late stat. | 250 | 6 | 0 | 6 | PFA |
| **(b)** | *A.b.* ATTC 17978 | 2.5 | Early stat. | 250 | 7 | 0 | 7 | PFA |
| | | 2.5 | Early stat. | 250 | 13 | 0 | 13 | PFA |
| **(c)** | *A.b.* ATTC 17978 | Harvested from plate | | 941 | 178 | not counted | | PFA |
| **(d)** | *A. baylyi* WT | Harvested from plate | | 2890 | 20 | not counted | | PFA |
| **(e)** | *A.b.* SC2151 | Harvested from plate | | 1184 | 95 | not counted | | PFA |
| **(f)** | *A.b.* SC1846 | Early stat. | | 747 | 0 | 0 | 0 | PFA |
| | | Harvested from plate | | 732 | 0 | 0 | 0 | PFA |

**Table S2: Antimicrobial resistance profiles of the parental strains and recombinant clones.** Inhibition zone diameters (mm) determined by disk diffusion according to EUCAST methodology. Disk contents (µg). h, heteroresistance.

| | **Inhibition zone diameters [mm]** | | | | | | | | |
|---|---|---|---|---|---|---|---|---|---|
| **Strain/ Clone** | **Mero-penem (10)** | **Imi-penem (10)** | **Cefta-zidime (30)** | **Pipera-cillin (30)** | **Pipera-cillin-Tazo-bactam (36)** | **Tige-cycline (15)** | **Trimethoprim/ Sulfamethoxazole (25)** | **Levo-floxacin (5)** | **Cipro-floxacin (5)** |
| **SC1846** | 6 | 9 | 13 | 6 | 6 | 17 | 6 | 9 | 6 |
| **SC2151** | 28 | 26 | 22 | 13 | 20 | 21 | 6 | 27 | 27 |
| **1** | 6 | 7 | 13 | 6 | 6 | 17 | 6 | 9 | 6 |
| **2** | 6 | 7 | 13 | 6 | 6 | 17 | 6 | 9 | 6 |
| **3** | 6 | 8 | 12 | 6 | 6 | 19 | 25[h] | 24 | 26 |
| **4+** | 6 | 7 | 14 | 6 | 6 | 20 | 6 | 12 | 6 |
| **5** | 6 | 7 | 13 | 6 | 6 | 17 | 6 | 9 | 6 |
| **6+** | 6 | 9 | 24 | 6 | 6 | 22 | 6 | 26 | 27 |
| **7-** | 6 | 13 | 13 | 6 | 6 | 18 | 6 | 9 | 6 |
| **8** | 6 | 7 | 13 | 6 | 6 | 17 | 6 | 9 | 6 |
| **9+** | 6 | 9 | 14 | 6 | 6 | 22 | 6 | 12 | 6 |
| **10-** | 6 | 8 | 14 | 6 | 6 | 21 | 6 | 6 | 6 |
| **11-** | 6 | 9 | 14 | 6 | 6 | 21 | 6 | 12 | 6 |
| **12+** | 6 | 7 | 14 | 6 | 6 | 18 | 6 | 9 | 6 |
| **13** | 6 | 8 | 23 | 6 | 6 | 17 | 6 | 21 | 24 |
| **14+** | 6 | 9 | 25 | 6 | 6 | 22 | 6 | 26 | 27 |
| **15** | 6 | 10 | 23 | 6 | 6 | 16 | 6 | 27 | 29 |
| **16** | 6 | 10 | 21 | 6 | 6 | 17 | 6 | 22 | 23 |
| **17-** | 6 | 9 | 8 | 6 | 6 | 17 | 6 | 6 | 6 |
| **18** | 6 | 9 | 14 | 6 | 6 | 21 | 6 | 12 | 6 |
| **19** | 6 | 10 | 25 | 6 | 6 | 22 | 6 | 26 | 26 |
| **20-** | 6 | 8 | 24 | 6 | 6 | 20 | 6 | 26 | 25 |
| **21-** | 6 | 7 | 24 | 6 | 6 | 20 | 6 | 25 | 26 |
| **22+** | 6 | 8 | 12 | 6 | 6 | 16 | 6 | 6 | 6 |

**Table S3: Impact of the recombination events on the gene sets of the recombinant clones.** PPanGGOLiN groups genes with >85% sequence similarity into gene families. The number of coding sequences (CDS) assigned to the lost/gained families is given in the corresponding columns.

| | SC1846 lost | | | | SC2151 acquired | | | | Net change | |
|---|---|---|---|---|---|---|---|---|---|---|
| Clone | Families | CDS | % Families | % CDS | Families | CDS | % Families | % CDS | Families | CDS |
| **Clone16** | 125 | 141 | 3.4 | 3.6 | 111 | 113 | 17.3 | 17.1 | -14 | -28 |
| **Clone10-** | 30 | 30 | 0.8 | 0.8 | 61 | 61 | 9.5 | 9.2 | 31 | 31 |
| **Clone11-** | 53 | 55 | 1.4 | 1.4 | 72 | 72 | 11.2 | 10.9 | 19 | 17 |
| **Clone13** | 93 | 109 | 2.5 | 2.8 | 118 | 119 | 18.4 | 18 | 25 | 10 |
| **Clone3** | 91 | 93 | 2.5 | 2.4 | 120 | 121 | 18.7 | 18.3 | 29 | 28 |
| **Clone18** | 11 | 11 | 0.3 | 0.3 | 12 | 12 | 1.9 | 1.8 | 1 | 1 |
| **Clone15** | 100 | 116 | 2.7 | 3 | 124 | 125 | 19.3 | 18.9 | 24 | 9 |
| **Clone22+** | 17 | 17 | 0.5 | 0.4 | 55 | 55 | 8.6 | 8.3 | 38 | 38 |
| **Clone21-** | 107 | 123 | 2.9 | 3.2 | 129 | 130 | 20.1 | 19.7 | 22 | 7 |
| **Clone20-** | 110 | 126 | 3 | 3.2 | 132 | 133 | 20.6 | 20.1 | 22 | 7 |
| **Clone14+** | 115 | 131 | 3.1 | 3.4 | 128 | 129 | 20 | 19.5 | 13 | -2 |
| **Clone6+** | 97 | 99 | 2.6 | 2.5 | 142 | 143 | 22.2 | 21.6 | 45 | 44 |
| **Clone19** | 112 | 128 | 3 | 3.3 | 133 | 134 | 20.7 | 20.3 | 21 | 6 |
| **Clone9+** | 45 | 45 | 1.2 | 1.2 | 63 | 63 | 9.8 | 9.5 | 18 | 18 |
| **Mean** | **79** | **87.4** | **2.1** | **2.3** | **100** | **100,7** | **15.6** | **15.2** | **21** | **13.3** |

**Table S4: Size (in bp) of chromosomal and plasmid sequences across original strains and recombinant clones.** Four distinct plasmids were identified (n.d. = not detected).

| Strain | Chromosome size [bp] | Plasmid-1 size [bp\| | Plasmid-2 size [bp\| | Plasmid-3 size [bp\| | Plasmid-4 size [bp\| |
|---|---|---|---|---|---|
| Clone10- | 4,160,971 | 17,284 | n.d. | n.d. | n.d. |
| Clone11- | 4,131,238 | 17,284 | 5,281 | n.d. | n.d. |
| Clone12+ | 4,115,944 | 17,284 | 5,281 | n.d. | n.d. |
| Clone17- | 4,118,611 | 17,284 | 5,281 | n.d. | n.d. |
| Clone18 | 4,119,783 | 17,284 | 5,281 | n.d. | n.d. |
| Clone22+ | 4,164,841 | 17,284 | 5,281 | n.d. | n.d. |
| Clone21- | 4,130,070 | 17,284 | 5,281 | n.d. | n.d. |
| Clone20- | 4,130,237 | 17,284 | 5,281 | n.d. | n.d. |
| Clone4+ | 4,117,725 | 17,284 | 5,281 | n.d. | n.d. |
| Clone14+ | 4,144,172 | 17,284 | 5,281 | n.d. | n.d. |
| Clone6+ | 4,151,383 | 17,284 | 5,281 | n.d. | n.d. |
| Clone7- | 4,117,725 | 17,284 | 5,281 | n.d. | n.d. |
| Clone19 | 4,134,171 | 17,284 | 5,281 | n.d. | n.d. |
| Clone9+ | 4,114,031 | 17,284 | 5,281 | n.d. | n.d. |
| Clone16 | 4,094,063 | 17,284 | 5,281 | n.d. | n.d. |
| Clone13 | 4,146,993 | 17,284 | 5,281 | n.d. | n.d. |
| Clone3 | 4,129,058 | 17,284 | 5,281 | n.d. | n.d. |
| Clone1 | 4,115,058 | 17,284 | 5,281 | n.d. | n.d. |
| Clone15 | 4,146,239 | 17,284 | 5,281 | n.d. | n.d. |
| Clone2 | 4,108,706 | 17,284 | 5,281 | n.d. | n.d. |
| Clone5 | 4,115,067 | 17,284 | 5,281 | n.d. | n.d. |
| Clone8 | 4,107,829 | 17,284 | 5,281 | n.d. | n.d. |
| SC1846 | 4,115,058 | 17,284 | 5,281 | n.d. | n.d. |
| SC2151 | 4,006,716 | n.d. | n.d. | 13,409 | 11,302 |

**Table S5: Bacterial isolates included in this study and corresponding BioProject and BioSample accession numbers.**

| Strain | Alternative ID | Strain Type | BioProject | BioSample |
|---|---|---|---|---|
| MS317 | | Clinical isolate | PRJNA1516054 | SAMN62596865 |
| SC2072 | | Clinical isolate | PRJNA1328187 | SAMN51307162 |
| SC1842 | BMBF_258 | Clinical isolate | PRJNA1328187 | SAMN51307163 |
| SC1833 | ABC141 | Clinical isolate | PRJNA1255660 | SAMN48152296 |
| SC2076 | | Clinical isolate | PRJNA1328187 | SAMN51307164 |
| 1372 | | Clinical isolate | PRJNA901493 | SAMN31716253 |
| 2778 | | Clinical isolate | PRJNA901493 | SAMN31716261 |
| SC2073 | | Clinical isolate | PRJNA1516054 | SAMN62596863 |
| SC2075 | | Clinical isolate | PRJNA1328187 | SAMN51307166 |
| SC1846 | | Clinical isolate | PRJNA1516054 | SAMN62596862 |
| SC2151 | | Clinical isolate | PRJNA1516054 | SAMN62596864 |
| Clone1 | | Laboratory-derived clone | PRJNA1516054 | SAMN62808447 |
| Clone2 | | Laboratory-derived clone | PRJNA1516054 | SAMN62808451 |
| Clone3 | | Laboratory-derived clone | PRJNA1516054 | SAMN62808452 |
| Clone4+ | | Laboratory-derived clone | PRJNA1516054 | SAMN62808453 |
| Clone5 | | Laboratory-derived clone | PRJNA1516054 | SAMN62808454 |
| Clone6+ | | Laboratory-derived clone | PRJNA1516054 | SAMN62808455 |
| Clone7- | | Laboratory-derived clone | PRJNA1516054 | SAMN62808456 |
| Clone8 | | Laboratory-derived clone | PRJNA1516054 | SAMN62808457 |
| Clone9+ | | Laboratory-derived clone | PRJNA1516054 | SAMN62808458 |
| Clone10- | | Laboratory-derived clone | PRJNA1516054 | SAMN62808437 |
| Clone11- | | Laboratory-derived clone | PRJNA1516054 | SAMN62808438 |
| Clone12+ | | Laboratory-derived clone | PRJNA1516054 | SAMN62808439 |
| Clone13 | | Laboratory-derived clone | PRJNA1516054 | SAMN62808440 |
| Clone14+ | | Laboratory-derived clone | PRJNA1516054 | SAMN62808441 |
| Clone15 | | Laboratory-derived clone | PRJNA1516054 | SAMN62808442 |
| Clone16 | | Laboratory-derived clone | PRJNA1516054 | SAMN62808443 |
| Clone17- | | Laboratory-derived clone | PRJNA1516054 | SAMN62808444 |
| Clone18 | | Laboratory-derived clone | PRJNA1516054 | SAMN62808445 |
| Clone19 | | Laboratory-derived clone | PRJNA1516054 | SAMN62808446 |

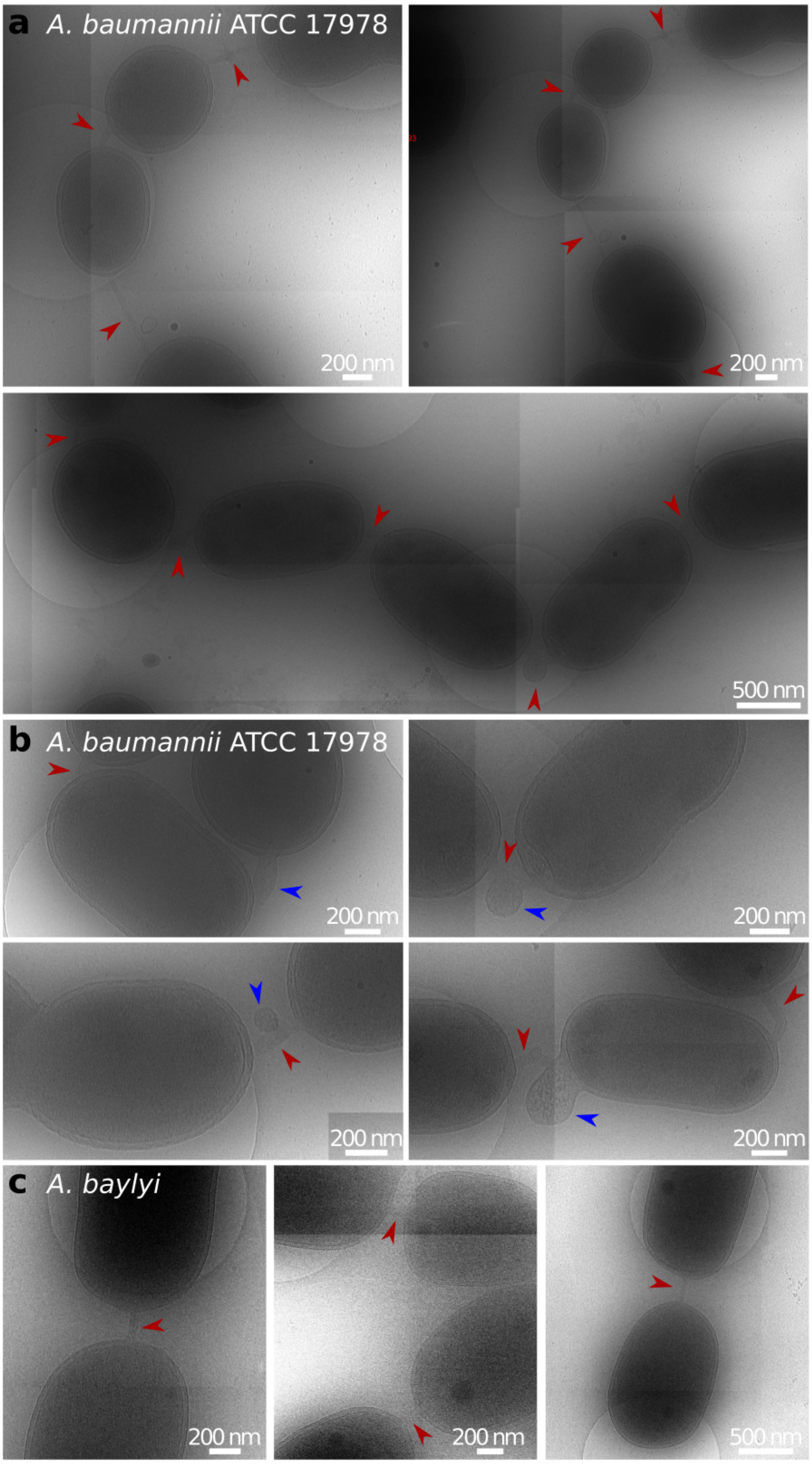


**Fig. S1: Cryo-electron micrographs of *A. baumannii* and *A. baylyi* strains.**
(a) Chains of *A.b.* ATCC 17978 cells connected by membrane conduits (indicated by red arrows).
(b) *A.b.* ATCC 17978 cells connected by conduits showing a thickening of the conduit diameter and lateral, electron-dense protrusion in the middle of the conduit or directly cell-adjacent (indicated by blue arrows).
(c) *A. baylyi* cells connected by membrane conduits (indicated by red arrows). A total of 20 membrane conduits were found in the 2890 *A. baylyi* cells examined (average frequency of 0.7%).

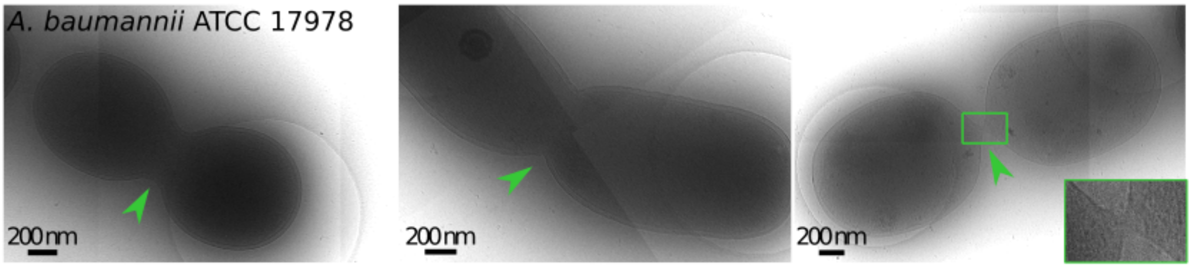


**Fig. S2: Exemplary cryo-electron micrographs of *A. baumannii* ATCC 17978 cell division septa (n>200).** Division septa are indicated by green arrows. In the inset, the position of the membrane connection is magnified. Early stages of cell division are characterized by deep invaginations of the cell envelope. In late-stage division intermediates, the daughter cells were nearly separated but remained connected by a narrow membrane bridge at the division site. Those bridges appeared distinct from membrane conduits, which were characterized by a relatively uniform diameter and the retention of the bacterial cell envelope.

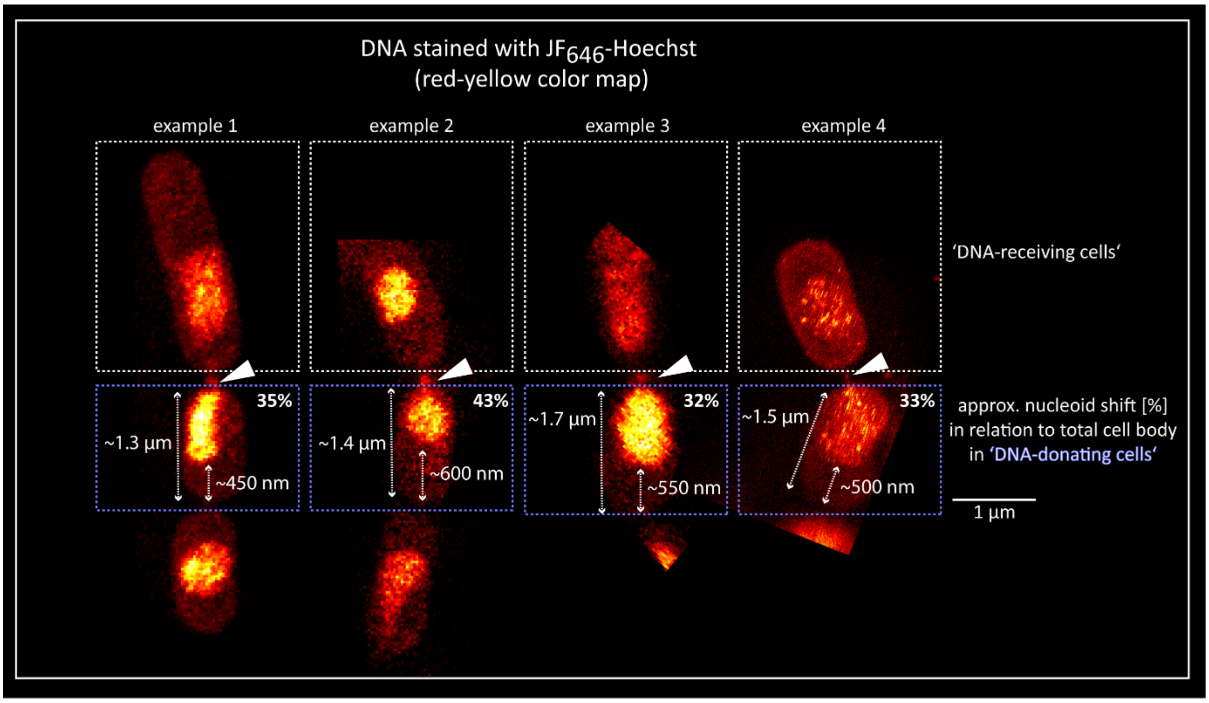


**Fig. S3: Approximate nucleoid shift in relation to the total cell body in *A. baumannii* ATCC 17978 developing conjugative bridges facilitating DNA transfer**. DNA stained with $JF_{646}$-Hoechst (red-yellow color map). Average shift = ~36%

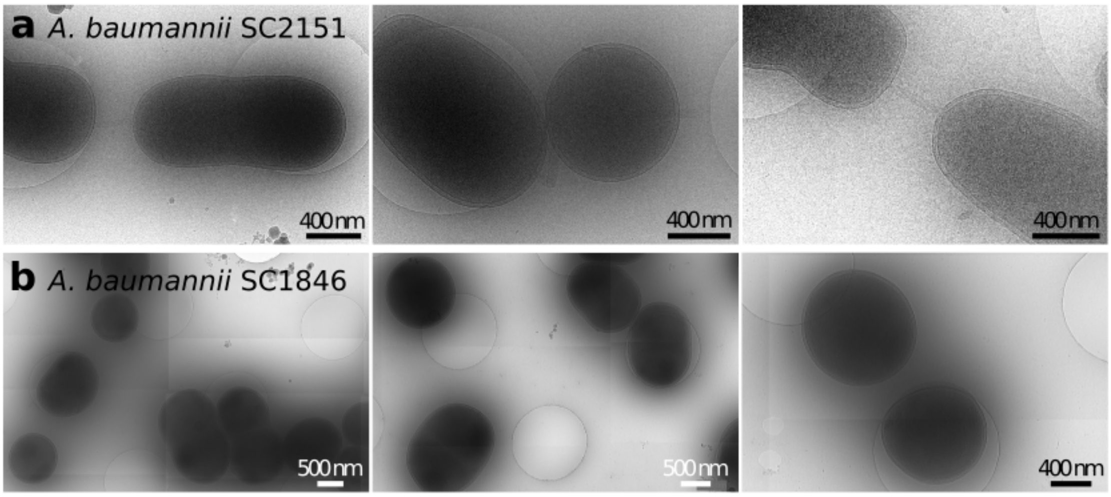


**Fig. S4: Cryo-electron micrographs of *A. baumannii* strains.**
**(a)** *A. baumannii* SC2151 cells connected by membrane conduits.
**(b)** *A. baumannii* SC1846 cells prepared under identical conditions and imaged optimized for conduit identification. No conduits were found in 747 cells grown in liquid culture and 732 cells harvested directly from plate.

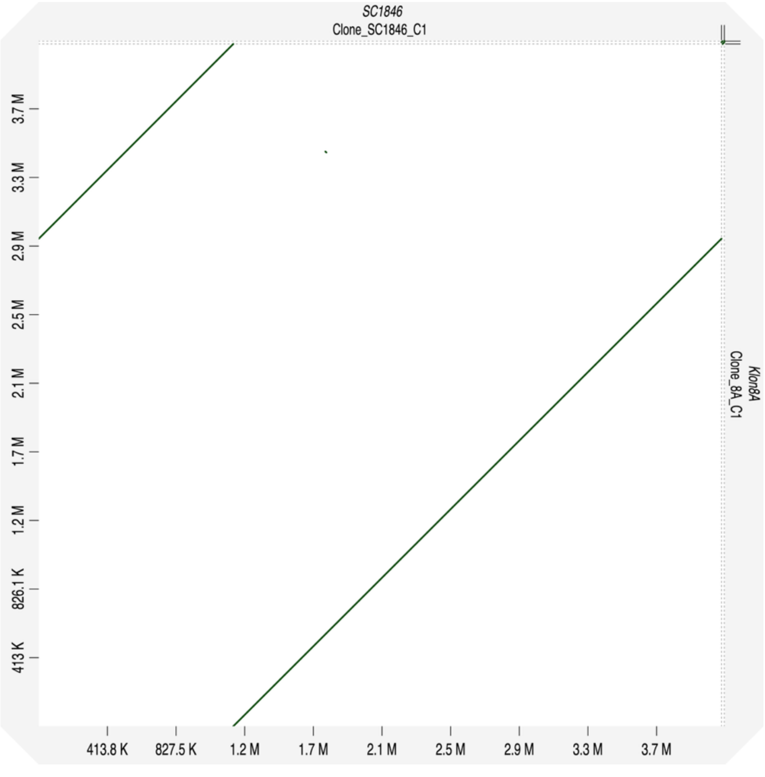


**Fig. S5. Dot plot of the bacterial chromosomes of *A. baumannii* SC1846 and one recombinant that acquired the resistance via point mutations.** The clone is representative for the group of eight recombinants with Mash distance of close to zero when compared to SC1846. Genomic positions are given as axis labels. Green diagonal lines represent regions of genomic sequence similarity and co-linearity.

```
Clone 5     pgap_000474  EMRTGEGKTLMGTLACYLNALSGEGVHVITVNDYLAQRDAELNRPLFEFL
Clone 8     pgap_000625  EMRTGEGKTLMGTLACYLNALSGEGVHVITVNDYLAQRDAELNRPLFEFL
Clone 4+    pgap_000861  EMRTGEGKTLMGTLACYLNALSGEGVHVITVNDYLAQRDAELNRPLFEFL
Clone 2     pgap_000003  EMRTGEGKTLMGTLACYLNALSGEGVHVITVNDYLAQRDAELNRPLFEFL
Clone 1     pgap_001380  EMRTGEGKTLMGTLACYLNALSGEGVHVITVNDYLAQHDAELNRPLFEFL
SC1846      pgap_001758  EMRTGEGKTLMGTLACYLNALSGEGVHVITVNDYLAQRDAELNRPLFEFL
Clone 7-    pgap_002397  EMRTGEGKTLMGTLACYLNALSGEGVHVITVNDYLAQRDAELNRPLFEFL
Clone 17-   pgap_002517  EMRTGEGKTLMGTLACYLNALSGEGVHVITVNDYLAQRDAELNRPLFEFL
Clone 12+   pgap_002664  EMRTGEGKTLMGTLACYLNALSGEGVHVITVNDYLAQRDAELNRPLFEFL
SC2151      pgap_001359  EMRTGEGKTLMDTLACYLNALSGEGVHVITVNDYLAQRDAELNRPLFEFL
                         ***********.**************************:***********

Clone 5     pgap_000474  HQAVEAKEGLAIQPENQTLATTTFQNYFRLYKKLSGMTGTADTEAAEMKE
Clone 8     pgap_000625  HQAVEAKEGLAIQPENQTLATTTFQNYFRLYKKLSGMTGTADTEAAEMKE
Clone 4+    pgap_000861  HQAVEAKEGLAIQPENQTLATTTFQNYFCLYKKLSGMTGTADTEAAEMKE
Clone 2     pgap_000003  HQAVEAKEGLAIQPENQTLATTTFQNYFRLYKKLSGMTGTADTEAAEMKE
Clone 1     pgap_001380  HQAVEAKEGLAIQPENQTLATTTFQNYFRLYKKLSGMTGTADTEAAEMKE
SC1846      pgap_001758  HQAVEAKEGLAIQPENQTLATTTFQNYFRLYKKLSGMTGTADTEAAEMKE
Clone 7-    pgap_002397  HQAVEAKEGLAIQPENQTLATTTFQNYFRLYKKLSGMTGTADTEAAEMKE
Clone 17-   pgap_002517  HQAVEAKEGLAIQPENQTLATTTFQNYFRLYKKLSGMTGTADTEAAEMKE
Clone 12+   pgap_002664  HQAVEAKEGLAIQPENQTLATTTFQNYFRLYKKLSGMTGTADTEAAEMKE
SC2151      pgap_001359  HQAVEAKEGLTIQPENQTLATTTFQNYFRLYKKLSGMTGTADTEAAEMKE
                         **********:***************** *********************

Clone 5     pgap_000474  LIGTATI EASEILSSKLKQAGIIMEVLNAKQIIEREADI IAQASSPNAVTI
Clone 8     pgap_000625  LIGTATI EASEILSSKLKQAGIIMEVLNAKQIIEREADI IAQASSPNAVTI
Clone 4+    pgap_000861  LIGTATI EASEILSSKLKQAGIIMEVLNAKQIIEREADI IAQAGSPNAVTI
Clone 2     pgap_000003  LIGTATI EASEILSSKLKQAGIIMEVLNAKQIIEREADI IAQAGSPNAVTI
Clone 1     pgap_001380  LIGTATI EASEILSSKLKQAGIIMEVLNAKQIIEREADI IAQAGSPNAVTI
SC1846      pgap_001758  LIGTATI EASEILSSKLKQAGIIMEVLNAKQIIEREADI IAQAGSPNAVTI
Clone 7-    pgap_002397  LIGTATI EASEILSSKLKQAGIIMEVLNAKQIIEREADI IAQAGSPNAVTI
Clone 17-   pgap_002517  LIGTATI EASEILSSKLKQAGIIMEVLNAKQIIEREADI IAQAGSPNAVTI
Clone 12+   pgap_002664  LIGTATI EASEILSSKLKQAGIIMEVLNAKQIIEREADI IAQAGSPNAVTI
SC2151      pgap_001359  LIGTATI EASEILSSKLKQAGIIMEVLNAKQIIEREADI IAQAGSPNAVTI
                         *********************************************.*******

Clone 5     pgap_000474  VVAMMRAMGLKEDEAIDIKMVSRSIENAQRKVEARNFDIRKNLLKYDDVN
Clone 8     pgap_000625  VVAMMRAMGLKEDEAIDIKMVSRSIENAQRKVEARNFDIRKNLLKYDDVN
Clone 4+    pgap_000861  VVAMMRAMGLKEDEAIDIKMVSRSIENAQRKVEARNFDIRKNLLKYDDVN
Clone 2     pgap_000003  VVAMMRAMGLKEDEAIDIKMVSRSIENAQRKVEARKFDIRKNLLKYDDVN
Clone 1     pgap_001380  VVAMMRAMGLKEDEAIDIKMVSRSIENAQRKVEARNFDIRKNLLKYDDVN
SC1846      pgap_001758  VVAMMRAMGLKEDEAIDIKMVSRSIENAQRKVEARNFDIRKNLLKYDDVN
Clone 7-    pgap_002397  VVAMMRAMGLKEDEAIDIKMVSRSIENAQRKVEARNFDIRKNLLKYDDVN
Clone 17-   pgap_002517  VVAMMRAMGLKEDEAIDIKMVSRSIENAQRKVEARNFDIRKNLLKYDDVN
Clone 12+   pgap_002664  VVAMMRAMGLKEDEAIDIKMVSRSIENAQRKVEARNFDIRKNLLKYDDVN
SC2151      pgap_001359  VVAMMRAMGLKEDEAIDIKMVSRSIENAQRKVEARNFDIRKNLLKYDDVN
                         *********************************** **************
```

**Fig. S6. Multiple sequence alignment of SecA from the 8 non-recombinant clones and the variant encoded in SC2151.** Only alignment blocks containing at least one variable position (red boxes) are shown. None of the clones carried the SC2151 variant of SecA, which strongly suggests that the observed point mutations occurred *de novo* in each clone.

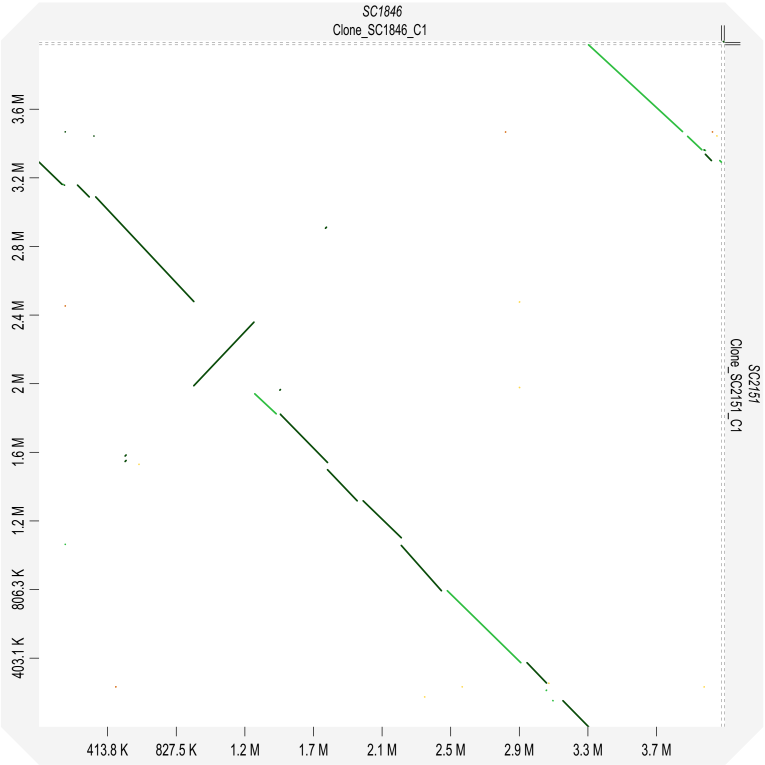


**Fig. S7. Dot plot of the bacterial chromosomes of *A. baumannii* SC1846 and *A. baumannii* SC2151.** Genomic positions are given as axis labels. Diagonal lines represent regions of genomic co-linearity. Change in line orientation indicates inversion.

```
Clone 22+   pgap_001639   EMRTGEGKTLMDTLACYLNALSGEGVHVITVNDYLAQRDAELNRPLFEFL
SC1846      pgap_001758   EMRTGEGKTLMGTLACYLNALSGEGVHVITVNDYLAQRDAELNRPLFEFL
Clone16     pgap_002950   EMRTGEGKTLMDTLACYLNALSGEGVHVITVNDYLAQRDAELNRPLFEFL
Clone10-    pgap_000057   EMRTGEGKTLMDTLACYLNALSGEGVHVITVNDYLAQRDAELNRPLFEFL
Clone 11-   pgap_001774   EMRTGEGKTLMDTLACYLNALSGEGVHVITVNDYLAQRDAELNRPLFEFL
Clone13     pgap_000047   EMRTGEGKTLMDTLACYLNALSGEGVHVITVNDYLAQRDAELNRPLFEFL
Clone3      pgap_000821   EMRTGEGKTLMDTLACYLNALSGEGVHVITVNDYLAQRDAELNRPLFEFL
Clone 18-   pgap_002438   EMRTGEGKTLMDTLACYLNALSGEGVHVITVNDYLAQRDAELNRPLFEFL
Clone 9+    pgap_002348   EMRTGEGKTLMDTLACYLNALSGEGVHVITVNDYLAQRDAELNRPLFEFL
SC2151      pgap_001359   EMRTGEGKTLMDTLACYLNALSGEGVHVITVNDYLAQRDAELNRPLFEFL
Clone 19    pgap_000645   EMRTGEGKTLMDTLACYLNALSGEGVHVITVNDYLAQRDAELNRPLFEFL
Clone 6+    pgap_002404   EMRTGEGKTLMDTLACYLNALSGEGVHVITVNDYLAQRDAELNRPLFEFL
Clone 14+   pgap_000087   EMRTGEGKTLMDTLACYLNALSGEGVHVITVNDYLAQRDAELNRPLFEFL
Clone 20-   pgap_002353   EMRTGEGKTLMDTLACYLNALSGEGVHVITVNDYLAQRDAELNRPLFEFL
Clone 21-   pgap_001800   EMRTGEGKTLMDTLACYLNALSGEGVHVITVNDYLAQRDAELNRPLFEFL
Clone 15    pgap_000087   EMRTGEGKTLMDTLACYLNALSGEGVHVITVNDYLAQRDAELNRPLFEFL
SC2150      pgap_002565   EMRTGEGKTLMGTLACYLNALSGEGVHVITVNDYLAQRDAELNRPLFEFL
                          ***********.**************************************

Clone 22+   pgap_001639   HQAVEAKEGLAIQPENQTLATTTFQNYFRLYKKLSGMTGTADTEAAEMKE
SC1846      pgap_001758   HQAVEAKEGLAIQPENQTLATTTFQNYFRLYKKLSGMTGTADTEAAEMKE
Clone16     pgap_002950   HQAVEAKEGLTIQPENQTLATTTFQNYFRLYKKLSGMTGTADTEAAEMKE
Clone10-    pgap_000057   HQAVEAKEGLTIQPENQTLATTTFQNYFRLYKKLSGMTGTADTEAAEMKE
Clone 11-   pgap_001774   HQAVEAKEGLTIQPENQTLATTTFQNYFRLYKKLSGMTGTADTEAAEMKE
Clone13     pgap_000047   HQAVEAKEGLTIQPENQTLATTTFQNYFRLYKKLSGMTGTADTEAAEMKE
Clone3      pgap_000821   HQAVEAKEGLTIQPENQTLATTTFQNYFRLYKKLSGMTGTADTEAAEMKE
Clone 18-   pgap_002438   HQAVEAKEGLTIQPENQTLATTTFQNYFRLYKKLSGMTGTADTEAAEMKE
Clone 9+    pgap_002348   HQAVEAKEGLTIQPENQTLATTTFQNYFRLYKKLSGMTGTADTEAAEMKE
SC2151      pgap_001359   HQAVEAKEGLTIQPENQTLATTTFQNYFRLYKKLSGMTGTADTEAAEMKE
Clone 19    pgap_000645   HQAVEAKEGLTIQPENQTLATTTFQNYFRLYKKLSGMTGTADTEAAEMKE
Clone 6+    pgap_002404   HQAVEAKEGLTIQPENQTLATTTFQNYFRLYKKLSGMTGTADTEAAEMKE
Clone 14+   pgap_000087   HQAVEAKEGLTIQPENQTLATTTFQNYFRLYKKLSGMTGTADTEAAEMKE
Clone 20-   pgap_002353   HQAVEAKEGLTIQPENQTLATTTFQNYFRLYKKLSGMTGTADTEAAEMKE
Clone 21-   pgap_001800   HQAVEAKEGLTIQPENQTLATTTFQNYFRLYKKLSGMTGTADTEAAEMKE
Clone 15    pgap_000087   HQAVEAKEGLTIQPENQTLATTTFQNYFRLYKKLSGMTGTADTEAAEMKE
SC2150      pgap_002565   HQAVEAKEGLTIQPENQTLATTTFQNYFRLYKKLSGMTGTADTEAAEMKE
                          **********:***************************************
```

**Fig. S8. Multiple sequence alignment of SecA from the 14 recombinant clones and the variant encoded in SC2150 ($NaN_3$-susceptible) and in SC2151.** Only alignment blocks containing at least one variable position (red boxes) between SC1846 and SC2151 are shown. $NaN_3$ resistance correlates exactly with the G->D substitution relative to SC1846 in the first alignment block.

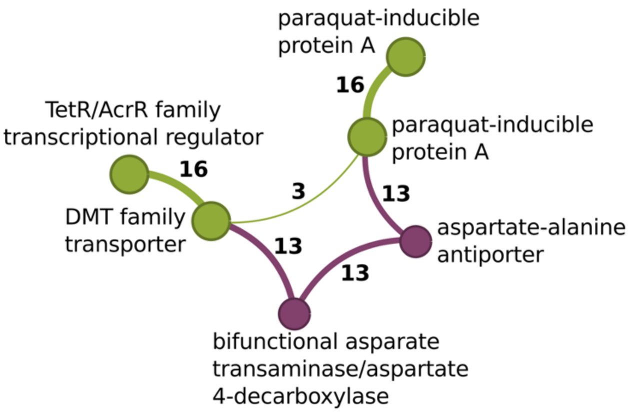


**Fig. S9. Recombination removes a functional unit of two genes involved in amino acid metabolism.** Nodes represent genes, and node diameter indicates the number of genomes harbouring the respective gene. Two nodes are connected by an edge if the corresponding genes occur next to each other in at least one of the analysed genomes. Edge thickness reflects the number of genomes in which the corresponding gene pair is adjacent; the number of genomes is indicated next to each edge. Green nodes and edges represent genes and gene adjacencies shared by the two parental strains and all 14 recombinant clones. In SC1846, a bifunctional aspartate transaminase/aspartate 4-decarboxylase and an aspartate-alanine antiporter occur next to each other and are flanked by a DMT family transporter and a paraquat-inducible protein A (violet path). In the corresponding region of SC2151, the two genes involved in amino acid metabolism are absent, leaving the two flanking genes as adjacent neighbours (yellow edge labelled 3). In two recombinant clones, recombination resulted in loss of the bifunctional aspartate transaminase/aspartate 4-decarboxylase and the aspartate-alanine antiporter.

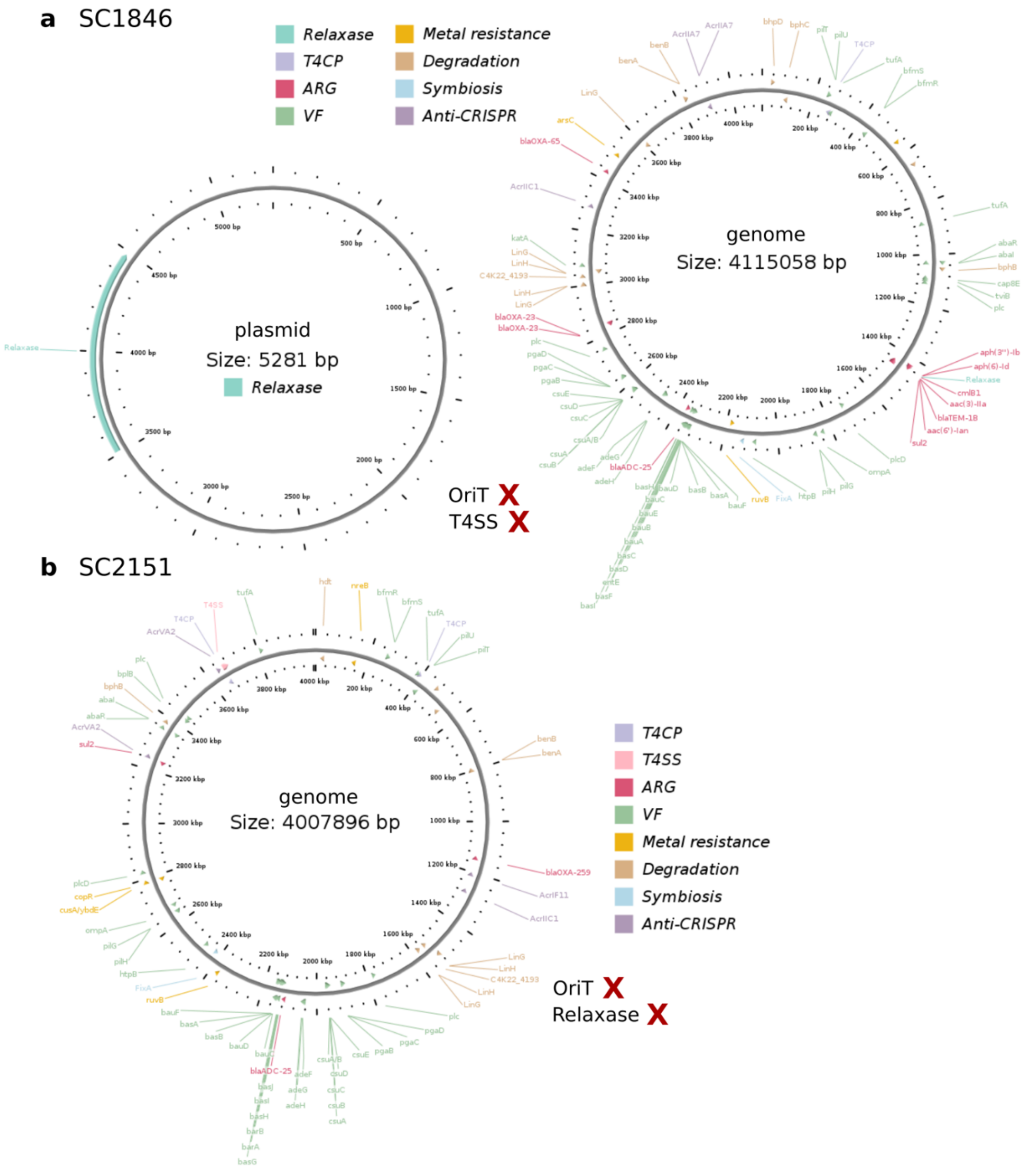


**Fig. S10. Prediction of conjugative genetic elements in different *A. baumannii* strains using oriTfinder, a web-based tool. The analysis predicts the origin of transfer (*oriT*), relaxase genes, type IV secretion system (T4SS), and type IV coupling protein (T4CP), as well as antibiotic resistance genes (ARGs) and virulence factor genes (VFs).**

(a) The *A. baumannii* SC1846 genome and plasmids are not predicted to contain an *oriT* or T4SS genes.

(b) The *A. baumannii* SC2151 genome is not predicted to contain an *oriT* or relaxase genes.
